# One lysine, two acyl marks: 2-hydroxyisobutyrylation and acetylation of G6PD differ in outcome and regulation

**DOI:** 10.64898/2026.10.29.762733

**Authors:** Santhosh Paramasivam, Omer Even Hen, Itamar Kass, Fang Wu, Eyal Arbely

## Abstract

Lysine 2-hydroxyisobutyrylation is a widespread post-translational modification enriched in enzymes of central carbon metabolism, yet its site-specific biochemical consequences remain poorly defined and usually inferred from amino acid substitutions or heterogeneously modified protein preparations. Using genetic code expansion technology, we generated homogeneous human glucose-6-phosphate dehydrogenase (G6PD) carrying an authentic 2-hydroxyisobutyryl group at nine individual lysine residues. 2-hydroxyisobutyrylation affected G6PD activity in a strongly site-dependent manner: modification within or adjacent to the catalytic pocket (K171, K205, K360), or at the tetramer interface (K288), abolished activity, whereas modification of K386, near the structural NADP^+^ binding site, reduced catalytic efficiency and destabilized the enzyme. HibK386 G6PD was less oligomeric, while inactive HibK288 G6PD formed stable oligomers, suggesting that in HibK288 G6PD, assembly and catalysis are separable. Molecular dynamics simulations indicated that 2-hydroxyisobutyrylation of K386 reorients the side chain toward the solvent and weakens hydrogen bonding and salt-bridge across the tetramer interface. K386 is also subject to acetylation, allowing the comparison between two acyl groups at one residue; acetylation destabilized G6PD without substantially impairing catalysis, whereas 2-hydroxyisobutyrylation did both, and the two marks were differentially recognized by their erasers. SIRT2 removed both and restored the activity of the purified enzyme, whereas SIRT1 readily deacetylated AcK386 but was a poor de-2-hydroxyisobutyrylase of this position. SIRT2 also de-2-hydroxyisobutyrylated HibK288 G6PD and PKM2 HibK62, but not PKM2 HibK115 or HibK305. Together, these findings identify site-specific 2-hydroxyisobutyrylation as a potent inhibitory modification of G6PD, demonstrate that chemically distinct acylations at the same site can exert different functional effects, and establish deacylase specificity as a function of both modification chemistry and residue context.

## Introduction

Lysine acylation, the covalent attachment of an acyl group to the ε-amine of lysine residues, is an abundant post-translational modification (PTM), with lysine acetylation being its most extensively studied form. Acylation of lysine side chains can occur enzymatically or non-enzymatically, and both routes require an acyl donor, such as an acyl-coenzyme A (acyl-CoA) thioester, thereby coupling acylation to intermediary metabolism^1^. Moreover, the occurrence of different acyl marks at the same lysine residue suggests an additional layer of regulation, in which the functional consequences of acylation may depend not only on the modified position but also on the identity of the acyl group. Advances in mass spectrometry (MS) have revealed that lysine 2-hydroxyisobutyrylation, the attachment of a 2-hydroxyisobutyryl (Hib) group (Supplementary Figure 1A), is widespread across bacteria, plants, and mammals^2–5^. It was first described as an active histone mark with a role in chromatin regulation^6^ and has since been mapped to thousands of non-histone sites across diverse organisms, with pronounced enrichment among enzymes of central carbon metabolism. Like acetylation, 2-hydroxyisobutyrylation neutralizes the positive charge of the lysine side chain, but the Hib group is bulkier and contains a hydroxyl group on a tertiary carbon that can act as both a hydrogen-bond (H-bond) donor and acceptor. At a given lysine, 2-hydroxyisobutyrylation can therefore replace an electrostatic interaction with a set of new H-bonding possibilities.

The cellular landscape of 2-hydroxyisobutyrylation is dynamically regulated by the opposing activities of lysine acyltransferases (‘writers’) and lysine deacylases (KDACs, ‘erasers’). In mammalian cells, p300, KAT7, KAT2A and Tip60 have been identified as lysine 2-hydroxyisobutyryltransferases, whereas de-2-hydroxyisobutyrylation can be catalyzed by the Zn^2+^-dependent histone deacetylases (HDACs) 1–3 and by the nicotinamide adenine dinucleotide (NAD^+^)-dependent sirtuins SIRT1, SIRT2, SIRT3, and SIRT7^5,7–11^. Since a given lysine residue can carry more than one type of acylation, each with distinct chemical characteristics, it is important to determine whether different acylations at the same position differentially affect protein function and whether KDACs can discriminate among the acyl groups occupying that position. Such selectivity would allow a cell to control the identity, rather than merely the level of acylation at an individual residue, yet this possibility has rarely been tested using full-length protein substrates with defined modification states.

Global proteomic studies have shown that lysine acylation, including 2-hydroxyisobutyrylation, is highly prevalent among metabolic enzymes, for which PTMs are an important regulatory strategy. In bacteria, for example, CobB removes Hib from enolase K343, thereby altering enzyme activity^12^. In human cells, p300-mediated 2-hydroxyisobutyrylation of enolase 1 (ENO1) at K281 modulates glycolytic flux in response to nutrient availability; unlike K228, which can also be acetylated, K281 was found to be only 2-hydroxyisobutyrylated^8^. Aldehyde dehydrogenase 1 family member A1 (ALDH1A1) is 2-hydroxyisobutyrylated at K260, where the modification promotes chaperone-mediated autophagic degradation; loss of this mark in bladder cancer stabilizes ALDH1A1 and is associated with poor prognosis^13^. Loss of SIRT3 likewise increases phosphofructokinase (PFK) 2-hydroxyisobutyrylation and shifts glycolytic flux during nephrogenesis^10^. These and other studies establish that 2-hydroxyisobutyrylation at a single site can govern the activity or stability of a metabolic enzyme and that KDACs play an important role in reversing this effect. However, the consequences of 2-hydroxyisobutyrylation were usually inferred from mutated proteins or potentially heterogeneously modified proteins.

Glucose-6-phosphate dehydrogenase (G6PD) occupies a critical metabolic branch point, where it can redirect carbon flux from glycolysis to the pentose phosphate pathway (PPP). As the rate-limiting enzyme of the PPP, G6PD controls the shunting of glucose-6-phosphate (G6P) from glycolysis to meet cellular demands for nucleotide precursors, reductive biosynthesis, and redox homeostasis^14,15^. Dysregulated G6PD activity is accordingly linked to various pathophysiological states, including cancer, infection, and clinical G6PD deficiency, the most common human enzymopathy, estimated to affect more than 400 million people worldwide^16–19^. Human G6PD functions as a homodimer or homotetramer, with evidence suggesting that the tetramer is the more active form^20,21^. Each G6PD protomer contains two distinct NADP^+^ binding sites: a catalytic site adjacent to the G6P-binding pocket and a distal structural site at which NADP^+^ binding stabilizes the active oligomeric state (Supplementary Figure 1B)^22–24^. G6PD activity is further modulated by various PTMs, including O-GlcNAcylation, phosphorylation, and acetylation^25–27^. For instance, acetylation at K89 promotes G6PD activity, whereas acetylation at K403, within the structural NADP^+^ binding site, completely suppresses G6PD activity^27–29^; the latter can be reversed by SIRT1-and SIRT2-mediated deacetylation^28,29^. Importantly, proteomic analyses across different species have found that G6PD is post-translationally modified by lysine 2-hydroxyisobutyrylation at multiple sites, some of which are also known acetylation sites^5,8,30–32^. However, the functional roles of these specific 2-hydroxyisobutyrylation events remain unknown.

Our current understanding of cellular events downstream of 2-hydroxyisobutyrylation and acetylation is derived mainly from mutational analyses. Although informative, the substituted residues (e.g., arginine or glutamine) differ from lysine or 2-hydroxyisobutyryl-lysine (HibK) in charge, volume, and hydrogen-bonding capacity and therefore neither faithfully mimic the structural and functional effects of lysine 2-hydroxyisobutyrylation nor reproduce the specific chemical properties that distinguish Hib from acetyl or lysine. Antibody-based readouts of endogenous modification further complicate interpretation since they may report on a heterogeneous mixture of modified and unmodified protein at unknown occupancy. These limitations leave several questions regarding site-specific lysine acylation unresolved. In particular, does G6PD 2-hydroxyisobutyrylation at a defined position have a regulatory role, and through what structural mechanism? When a single lysine in G6PD can carry either an acetyl or a Hib group, do the two marks produce the same functional outcome? Assuming a deacylase can remove both marks, would it discriminate between the two acyl modifications on the same residue?

In this study, we addressed these questions using genetic code expansion (GCE) technology to eliminate reliance on mimetic mutations and heterogeneous populations. Genetic code expansion employs an evolved aminoacyl-tRNA synthetase to charge its cognate tRNA with a non-canonical amino acid (ncAA) such as HibK or acetylated lysine (AcK). The ncAA is then incorporated cotranslationally in response to a unique codon, usually an in-frame UAG codon, placed at essentially any position in the protein of interest^33–35^. This approach yields homogeneous, quantitatively modified full-length protein carrying the authentic acyl group at a single, predetermined site. Using GCE technology, we studied the functional consequences of site-specific 2-hydroxyisobutyrylation of G6PD and the underlying regulatory mechanism. We identified 2-hydroxyisobutyrylation sites at which the modification completely inhibited enzymatic activity and found that 2-hydroxyisobutyrylation of K386 had a negative effect on both the activity and stability of G6PD, which we attributed to the loss of stabilizing interactions across the tetramer interface. In contrast, 2-hydroxyisobutyrylation of K288 inhibited the enzyme while increasing, rather than decreasing, the proportion of higher-order oligomers. Residue K386 can be modified by both acetylation and 2-hydroxyisobutyrylation, yet only the latter was inhibitory. We further found that SIRT1 preferentially targeted K386-acetylated G6PD over the 2-hydroxyisobutyrylated protein, whereas SIRT2 functioned as an efficient lysine deacylase capable of removing both acetylation and 2-hydroxyisobutyrylation in a selective manner. Specifically, SIRT2-mediated removal of Hib from K386 restored purified G6PD activity.

## Results

### Genetic code expansion enables site-specific 2-hydroxyisobutyrylation of nine lysine residues in human G6PD

Recent MS-based proteomic studies identified multiple lysine residues in human G6PD as 2-hydroxyisobutyrylation sites^5,8,30–32^. Nine of these residues were selected for the current study through the following prioritization process (Figure 1A). 2-hydroxyisobutyrylation sites in G6PD orthologs were retrieved from published MS data, and multiple sequence alignment was used to identify evolutionarily conserved modified lysine residues (Supplementary Figure 2). Superimposition of human G6PD structure with predicted G6PD structures was used to estimate the structural conservation of these sites (Supplementary Figure 3). The selection was further refined according to the position of each 2-hydroxyisobutyrylation site (e.g., near the active site or at subunit interfaces). Modified lysines close to the C-terminus were excluded because premature termination may yield potentially active C-truncated G6PD^29^. On this basis, we selected nine 2-hydroxyisobutyrylation sites: K82, K95, K97, K171, K205, K288, K360, K386, and K432 (Figure 1A). Residues K171, K205, and K360 are positioned near or within the catalytic pocket, and K288 is located at the tetramer interface. Residue K386 is part of an extended loop close to the structural NADP^+^ binding site and the dimer interface^22,23^; notably, the clinical G6PD Iowa variant (K386E) causes severe enzyme deficiency by disrupting NADP^+^ binding^36^, and the K386Q mutation significantly reduced G6PD activity^27^. Residues K171, K205, K288, and K360 are evolutionarily conserved (Supplementary Figure 2). Residues K82, K97, K171, K205, K360, K386, and K432 were identified as 2-hydroxyisobutyrylated in at least two different species, and residues K95, K97, K171, K386, and K432 are also known acetylation sites^37,38^, suggesting potential competition between acetylation and 2-hydroxyisobutyrylation.

**Figure 1.**
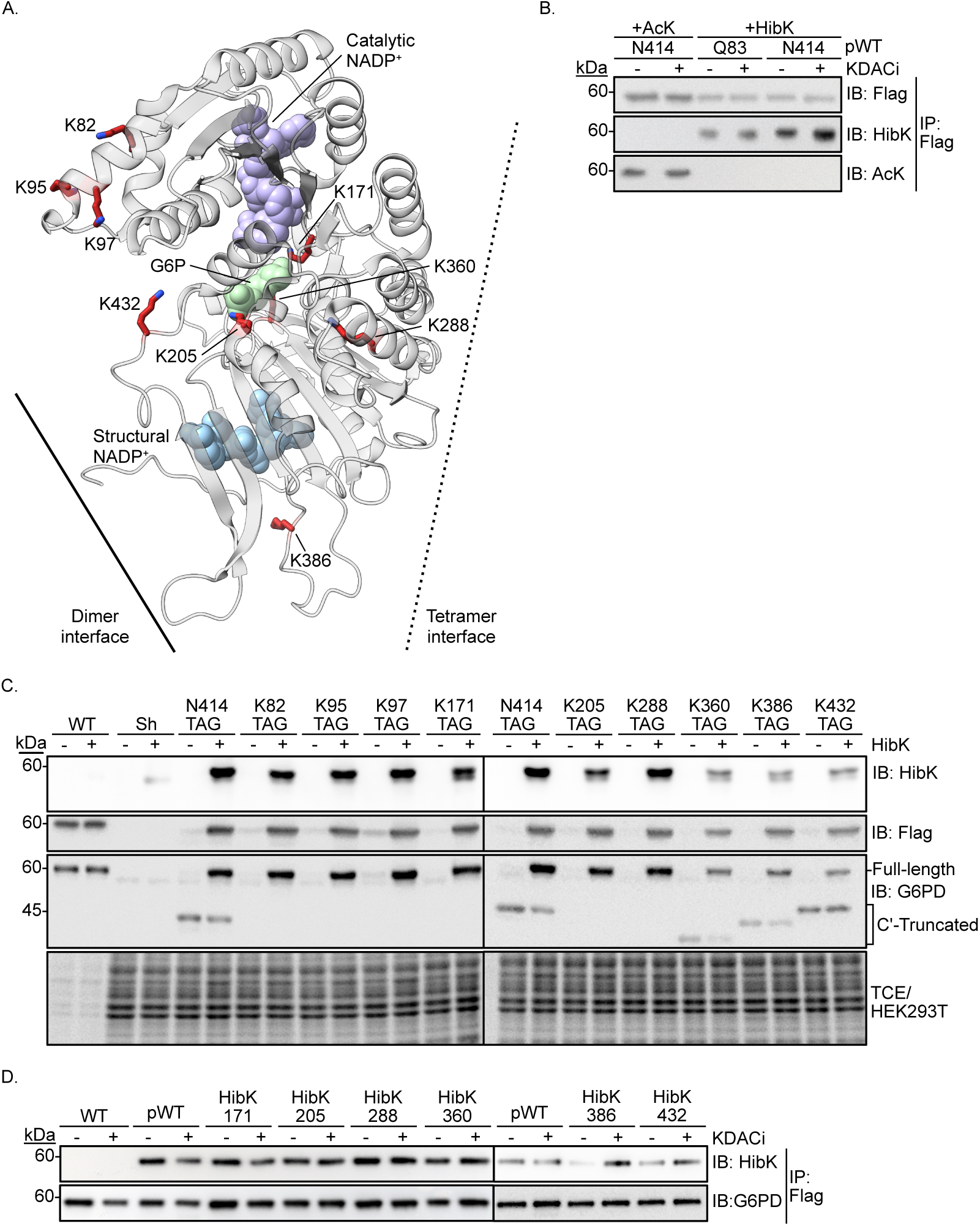
G6PD 2-hydroxyisobutyrylation sites, selection of pWT and expression of G6PD variants. **A.** Structure of inactive D200N human G6PD monomer (PDB ID: 7SNI) ^23^. The nine lysine 2-hydroxyisobutyrylation sites selected for this study are shown as sticks and highlighted in red. The dimer and tetramer interfaces are marked schematically by a continuous and a dashed line, respectively. Structural NADP^+^ and the substrates G6P and NADP^+^ are also shown. **B.** Validation of HibK414 G6PD as a non-2-hydroxyisobutyrylated control. Immunoblots show the modification level (anti-HibK or anti-AcK) of Flag-tagged G6PD variants expressed in cultured HEK293T cells. Immunoblot intensities of N414(HibK414) and Q83(HibK83) are displayed relative to the published control (AcK414) ^29^. **C.** HibK-dependent incorporation and expression of G6PD variants in mammalian cells. Cleared total lysates of HEK293T cells expressing the indicated G6PD variant, or the shRNA alone (sh) as a control, and cultured in the absence (-) or presence (+) of HibK, were resolved by SDS-PAGE. Immunoblots show 2-hydroxyisobutyrylation levels (anti-HibK), expression of full-length G6PD-Flag (anti-Flag), and expression of full-length and C-truncated G6PD (anti-G6PD). **D.** De-2-hydroxyisobutyrylation of G6PD variants by endogenous KDACs. Flag-tagged G6PD was immunoprecipitated from HEK293T cells expressing WT, pWT, or the indicated 2-hydroxyisobutyrylated variant, cultured without (-) or with (+) KDACi. Immunoblots show 2-hydroxyisobutyrylation levels (anti-HibK) and the amount of full-length G6PD loaded in each lane (anti-Flag).

To express site-specifically 2-hydroxyisobutyrylated G6PD in cultured mammalian cells, we established an expression system based on GCE technology^34^, using a single vector that carries all the required genetic components (Supplementary Figure 4A)^39^. HibK incorporation efficiency was evaluated with a dual-fluorescent reporter in which N-terminal mCherry is separated from C-terminal GFP by an in-frame TAG mutation (mCherry-TAG-GFP). Therefore, GFP expression depends on suppression of the UAG stop codon and incorporation of HibK, and the GFP/mCherry fluorescence ratio reports on incorporation efficiency. In cleared lysates of HEK293T cells expressing this reporter and cultured in media supplemented with 0 to 4 mM HibK, the GFP/mCherry ratio increased in a dose-dependent manner and plateaued at *∼*2 mM HibK (Supplementary Figure 4B), which was therefore selected as the optimal HibK concentration.

Measuring G6PD activity as a function of site-specific 2-hydroxyisobutyrylation requires a control for the non-acylated state. We therefore included two pseudo-wild-type (pWT) versions of G6PD, in which HibK was incorporated at a permissive site (Q83 or N414)^29^. The use of pWT G6PD ensures that the non-2-hydroxyisobutyrylated control is expressed at the same level as the studied 2-hydroxyisobutyrylated protein, and also accounts for potential unknown physiological effects that may arise from the amber-suppression machinery or the expression of truncated proteins. HibK incorporation into pWT G6PD was verified by Western blotting (Figure 1B), and, as previously established for the pWT used to study acetylated G6PD^29^, pWT 2-hydroxyisobutyrylation was independent of treatment with the KDAC inhibitors (KDACi) nicotinamide (NAM, a sirtuin inhibitor) and suberoylanilide hydroxamic acid (SAHA, an inhibitor of Zn^2+^-dependent HDACs) (Figure 1B). In addition, no cross-reactivity was found between anti-AcK and anti-HibK antibodies. For the current work, we chose HibK414 G6PD as the non-2-hydroxyisobutyrylated control (hereafter, pWT).

Next, pWT and the nine mono-2-hydroxyisobutyrylated, C-terminally Flag-tagged G6PD variants were expressed in HEK293T cells in which endogenous G6PD was knocked down. Immunoblot analysis with an antibody against the C-terminal Flag tag confirmed HibK-dependent expression of all nine full-length variants, with only minor variability in expression levels (Figure 1C). The anti-HibK antibody recognized the 2-hydroxyisobutyrylated variants but not WT G6PD, confirming that HibK incorporation depends on UAG suppression. pWT G6PD was expressed at levels comparable to those of the nine physiologically relevant 2-hydroxyisobutyrylated variants, whereas WT G6PD was expressed at much higher levels, as expected. In variants carrying the TAG mutation closer to the C-terminus (K360, K386, and K432), C-terminally truncated G6PD was detectable by Western blotting (Figure 1C). To test whether the 2-hydroxyisobutyrylated variants are de-2-hydroxyisobutyrylated by endogenous deacylases, cells were cultured without or with KDACi, Flag-tagged G6PD was immunoprecipitated, and 2-hydroxyisobutyrylation levels were monitored with an anti-HibK antibody (Figure 1D). KDACi treatment had a notable effect on 2-hydroxyisobutyrylation levels at K386 and, to a lesser extent, at K432. Together, these results establish a system for the robust expression of site-specifically 2-hydroxyisobutyrylated G6PD in mammalian cells for subsequent biochemical and functional experiments, and indicate that 2-hydroxyisobutyrylation of K386 is sensitive to endogenous KDAC activity.

### 2-hydroxyisobutyrylation inhibits G6PD activity in a position-dependent manner, with distinct effects on oligomerization

G6PD enzymatic activity was measured in cleared cell lysates using a continuous enzyme-coupled assay^40^. The activity of pWT G6PD was markedly higher than the background activity measured in lysates of shRNA-expressing cells (Supplementary Figure 4C). In addition, the catalytic activity of pWT G6PD was independent of KDAC inhibition and comparable to that of the previously established pWT (AcK414 G6PD)^29^, validating its use as a non-2-hydroxyisobutyrylated control in catalytic activity measurements (Supplementary Figure 4D). Finally, the activity measured in lysates of cells expressing G6PD TAG mutants and cultured without HibK, and therefore expressing C-truncated G6PD, was similar to background levels, indicating that the truncated proteins are inactive (Supplementary Figure 4C). Together, these controls confirm that the G6PD activity measured in cell lysates reports on exogenously expressed, site-specifically 2-hydroxyisobutyrylated G6PD.

The nine 2-hydroxyisobutyrylated G6PD variants were then expressed in HEK293T cells cultured in the absence or presence of KDACi. Enzymatic activity measured in cleared lysates showed that HibK82, HibK95, HibK97, and HibK432 G6PD were as active as the pWT control (Figure 2A and 2B), indicating that 2-hydroxyisobutyrylation at these positions does not substantively affect G6PD activity. In contrast, 2-hydroxyisobutyrylation at K171, K205, and K360, which are located within or adjacent to the catalytic pocket, essentially abolished activity, irrespective of KDACi treatment. 2-hydroxyisobutyrylation of K288, located at the tetramer interface^22,23^, likewise inhibited the enzyme both in the absence and in the presence of KDACi. Intriguingly, 2-hydroxyisobutyrylation of K386, which lies close to the structural NADP^+^ binding site, significantly reduced G6PD activity only upon KDACi treatment (Figure 2B), in agreement with the de-2-hydroxyisobutyrylation of HibK386 by endogenous KDACs detected by Western blotting (Figure 1D). Together, these data show that 2-hydroxyisobutyrylation of K171, K205, K288, K360, and K386 inhibits the activity of G6PD.

**Figure 2.**
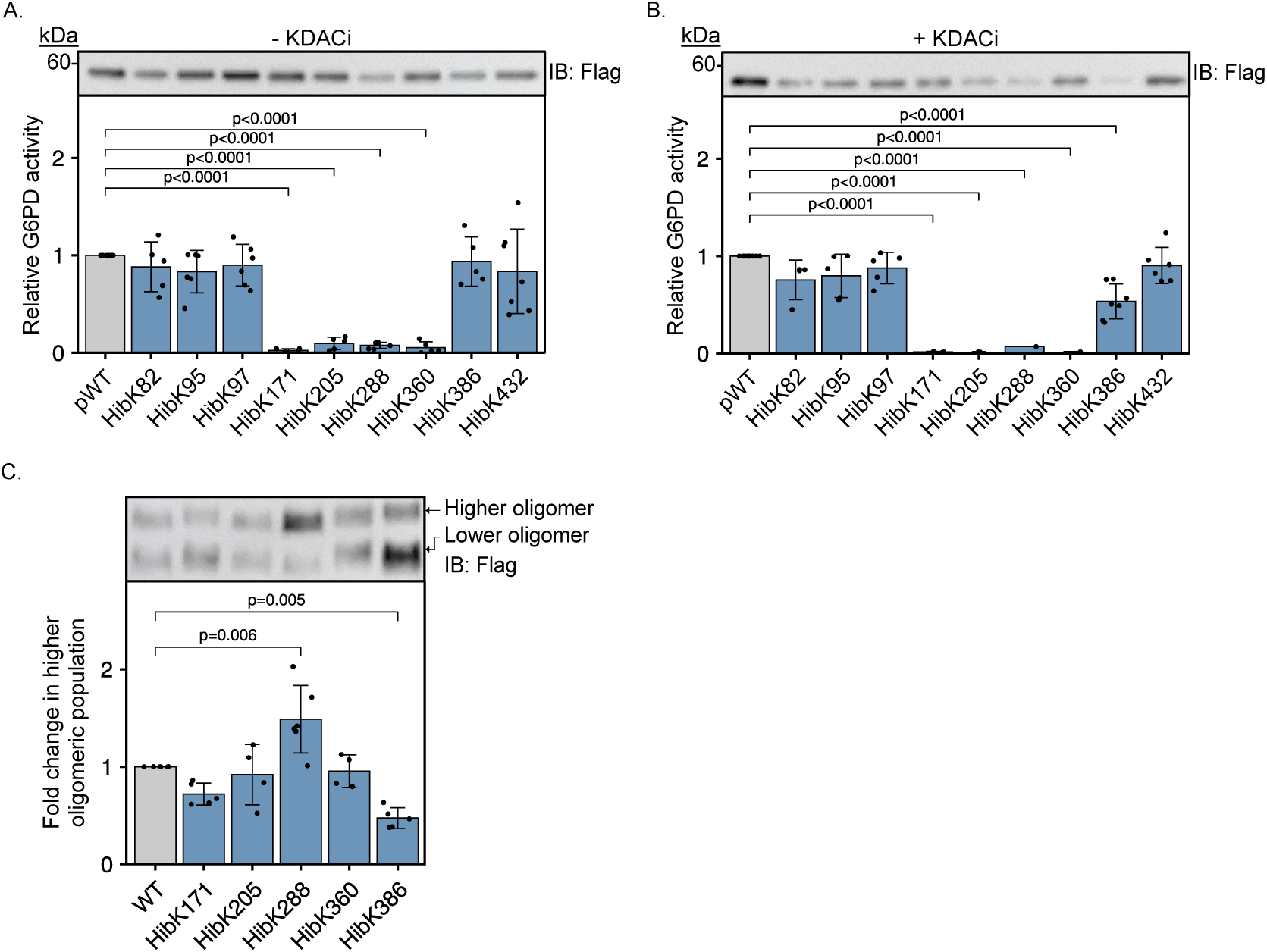
Regulation of G6PD activity and oligomerization by 2-hydroxyisobutyrylation. **A., B.** Activity of G6PD-Flag variants expressed in HEK293T cells cultured in the absence (A) or presence (B) of KDACi. Bars represent the *V*_max_ of exogenous G6PD measured in cleared cell lysates, normalized to immunoblot intensities of G6PD expression and displayed relative to pWT. **C.** Native PAGE analysis of G6PD variants expressed in HEK293T cells. Bars represent the ratio between the higher-order oligomeric species and total G6PD, determined by immunoblot quantification. Data are the mean *±* SD; *n ≥* 4 biological replicates. Statistical analysis was performed using one-way ANOVA followed by Tukey’s post hoc test.

Oligomerization is essential for G6PD activity, and many of the natural mutations that cause severe G6PD deficiency cluster around the dimer interface and the structural NADP^+^ binding site, compromising both enzyme stability and function^20,41^. To determine whether reduced oligomerization underlies the 2-hydroxyisobutyrylation-dependent loss of activity, we used native PAGE to evaluate the oligomeric state of the five inhibited G6PD variants (Figure 2C). 2-hydroxyisobutyrylation of K171, K205, or K360, which reside within the catalytic pocket, had no significant effect on the oligomeric state relative to the WT protein. In contrast, HibK288 G6PD displayed a greater tendency to form stable oligomers, even though 2-hydroxyisobutyrylation at this position significantly reduced enzyme activity. Importantly, HibK386 G6PD was less oligomeric than WT G6PD, in accordance with its lower enzymatic activity (Figure 2B) and with the position of K386 close to the structural NADP^+^ binding site and the dimer interface (Supplementary Figure 1B).

Taken together, these data indicate that site-specific 2-hydroxyisobutyrylation can inhibit G6PD activity, particularly at lysine residues within or close to the active site, and that 2-hydroxyisobutyrylation of K386, a substrate of endogenous KDACs, impairs both the activity and the oligomerization of G6PD.

### 2-hydroxyisobutyrylation of K386 destabilizes G6PD and impairs catalysis

Given the effects of K386 2-hydroxyisobutyrylation on G6PD oligomerization and activity, its KDAC-dependent regulation, and the identification of K386 as a known acetylation site, we focused our subsequent analyses on this residue. To assess the impact of K386 acylation on G6PD thermal stability, we measured the residual activity of WT, HibK386, and AcK386 G6PD in mammalian cell lysates following incubation over a range of increasing temperatures (Figure 3A). The measured *T*_1_*_/_*_2_ of WT G6PD was 47.0 *±* 0.2°C. In contrast, *T*_1_*_/_*_2_ values for the acylated variants could not be reliably determined because their thermal inactivation profiles lacked a detectable pretransition baseline. These results are consistent with a marked reduction in G6PD thermal stability upon acetylation or 2-hydroxyisobutyrylation of K386.

**Figure 3.**
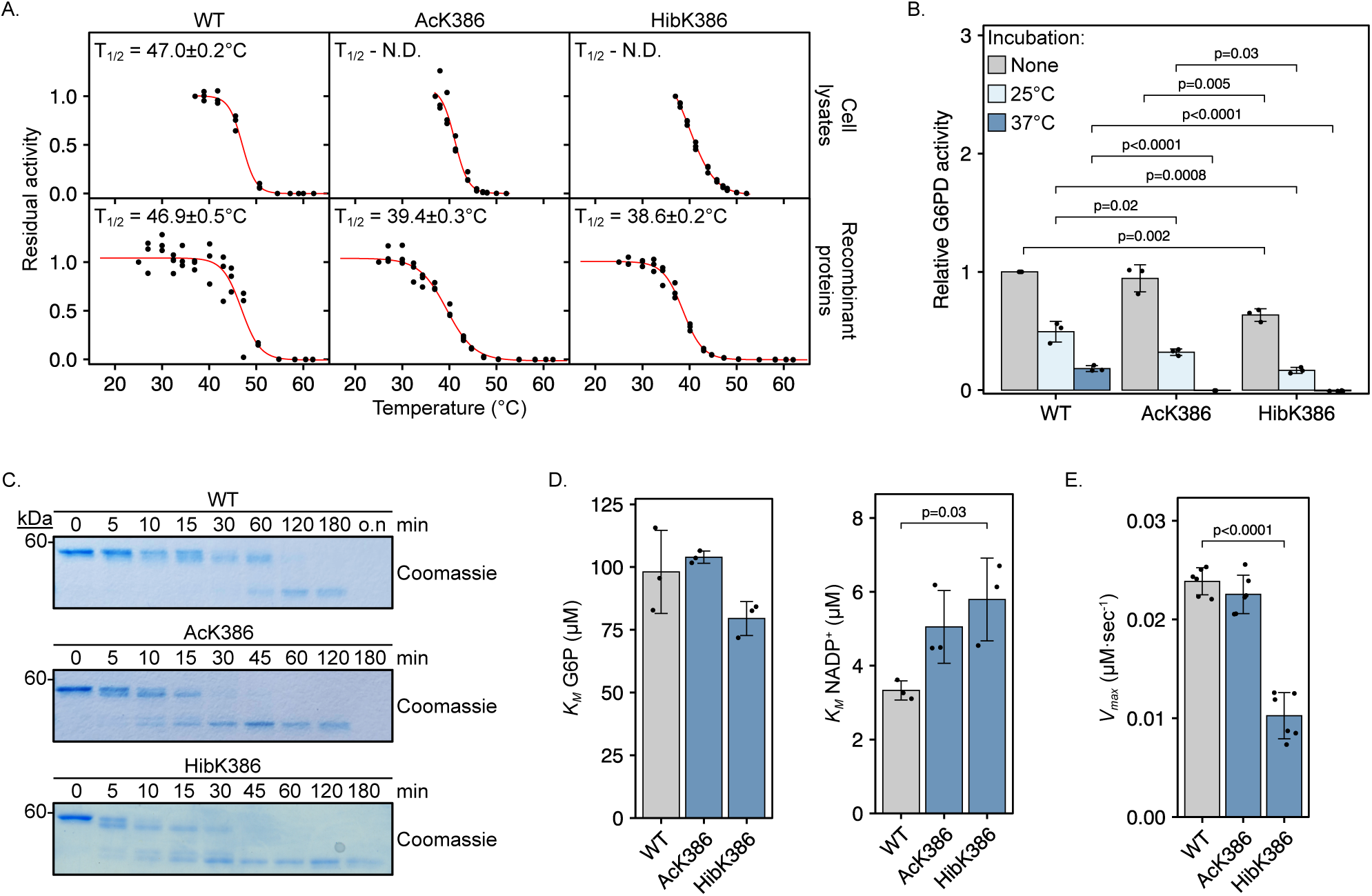
Acylation of K386 modulates G6PD stability and enzyme kinetics. **A.** Residual activity of WT and K386-modified G6PD after heat inactivation. The indicated G6PD variants, expressed in cultured HEK293T cells (top) or expressed recombinantly in bacteria and purified (bottom), were incubated at increasing temperatures before measurement of G6PD activity. Graphs show the residual activity as a function of incubation temperature, together with the fit of the data to a four-parameter logistic function (red line) from which *T*_1_*_/_*_2_ values were calculated. *T*_1_*_/_*_2_ values are reported *±* fitting error; *n* = 3. N.D., not determined. **B.** Residual activity of G6PD variants at physiologically relevant temperatures. Bars represent the activity of the indicated purified G6PD variants before and after overnight incubation at 25°C or 37°C, normalized to the activity of WT G6PD prior to incubation. Data are the mean *±* SD; *n* = 3. **C.** Limited proteolysis as a measure of conformational stability. Purified G6PD variants were incubated with chymotrypsin for the indicated times, and proteolytic degradation was monitored by SDS-PAGE and Coomassie staining. The persistence of the intact protein over time was used as a measure of resistance to proteolysis and, consequently, of conformational stability. **D.** Apparent Michaelis constants (*K*_M_^app^) for G6P and NADP^+^. The steady-state kinetics of WT and K386-modified G6PD were measured as a function of G6P or NADP concentration, and initial rates were fitted to the Michaelis–Menten equation, from which *K*_M_^app^ values were extracted. Bars represent the mean *±* SD; *n* = 3 independent measurements. **E.** Apparent maximal velocities (*V*_max_^app^) of WT and K386-modified G6PD. Steady-state kinetics were measured and *V*_max_^app^ values were extracted as described in panel D. Bars represent the mean *±* SD; *n* = 6 independent measurements. Statistical analysis was performed using one-way ANOVA followed by Tukey’s post hoc test (B, D & E).

To confirm these observations in a defined environment and to facilitate the biochemical characterization of WT and modified G6PD, we expressed WT, HibK386, and AcK386 G6PD in *E. coli*. Culture conditions were first optimized to maximize protein yield (Supplementary Figure 5A–C). The proteins were purified to homogeneity, and site-specific incorporation of the acylated lysine residue was confirmed by immunoblotting with anti-acetyllysine or anti-HibK antibodies and by LC-MS/MS (Supplementary Figure 5D and 5E). Residual activity measurements over a broader range of temperatures found that the *T*_1_*_/_*_2_ of purified WT G6PD was 46.9 *±* 0.5°C (Figure 3A), in agreement with measurements in cleared lysates. Importantly, both HibK386 and AcK386 variants exhibited a *T*_1_*_/_*_2_ of approximately 6–7°C lower than that of WT G6PD, indicating a decrease in protein stability following K386 2-hydroxyisobutyrylation or acetylation.

The thermal stability of purified WT and K386-acylated G6PD was further assessed by measuring enzyme activity before and after incubation at 25 or 37°C for *∼*20 h (Figure 3B). Acetylation of K386 had no effect on G6PD activity when measured prior to incubation, whereas the activity of HibK386 G6PD was significantly lower, reaching only *∼*60% of the WT level, in agreement with the activity measurements in cell lysates. Incubation at 25°C negatively affected the residual activity of all three proteins, with a stronger impact on AcK386 G6PD and a more pronounced impact on HibK386 G6PD. The difference was most distinct after incubation at 37°C, where both AcK386 and HibK386 G6PD lost catalytic activity completely, while WT G6PD retained *∼*15% of the activity measured before incubation (Figure 3B). Finally, we monitored the time-dependent in vitro digestion of recombinant WT and K386-acylated G6PD by chymotrypsin (Figure 3C). Separation of the digestion products by gel electrophoresis followed by Coomassie staining showed that both AcK386 and HibK386 G6PD were more susceptible to chymotrypsin digestion than WT G6PD, indicating lower conformational stability of K386-modified G6PD. Collectively, these data show that acylation of K386 compromises G6PD stability; in particular, K386 2-hydroxyisobutyrylation markedly reduced G6PD oligomerization, with clear indications of impaired thermal stability and reduced enzymatic activity.

To gain insight into the effect of K386 acylation on G6PD kinetics, we measured the steady-state kinetics of the recombinant enzymes. Initial velocities were measured as a function of G6P or NADP^+^ concentration while the other substrate was fixed at a saturating concentration, and apparent *V*_max_ and *K*_M_ values were calculated by fitting the data to the Michaelis–Menten equation (Supplementary Figure 6). The kinetic parameters of AcK386 G6PD were largely similar to those of the WT protein, with a notable decrease in catalytic efficiency (*k*_cat_/*K*_M_) for NADP^+^ (Figure 3D and 3E, and Supplementary Table 2). In contrast, 2-hydroxyisobutyrylation of K386 had a significant impact on G6PD kinetics with *∼*50% drop in *V*_max_. With G6P as the varied substrate, HibK386 G6PD maintained a *K*_M_ comparable to that of the WT protein, yet its catalytic efficiency was reduced by *∼*50%, as a consequence of a decreased *k*_cat_. With NADP^+^ as the varied substrate, HibK386 G6PD exhibited a *∼*1.8-fold increase in *K*_M_ and a *∼*50% decrease in *k*_cat_, together resulting in an *∼*75% decrease in *k*_cat_/*K*_M_.

Taken together, these results show that 2-hydroxyisobutyrylation of K386 has a greater functional impact than acetylation at the same residue, reducing both catalytic activity and oligomer stability.

### 2-hydroxyisobutyrylation of K386 remodels its local environment and weakens polar interactions across the tetramer interface

To elucidate the structural basis of the destabilization and inhibition caused by K386 2-hydroxyisobutyrylation, we first attempted to determine the crystal structure of HibK386 G6PD by X-ray crystallography. In the absence of crystals suitable for diffraction analysis, we performed molecular dynamics (MD) simulations of tetrameric WT and HibK386 G6PD bound to structural NADP^+^. Three independent, randomly seeded simulations of 490 ns each were performed for each variant, and the convergence of the root mean square deviation (RMSD) and radius of gyration (Rg) values was confirmed post-simulation (Supplementary Figure 7A and 7B). The average Cα RMSD was 0.39 *±* 0.07 nm for the WT tetramer and 0.25 *±* 0.04 nm for the HibK386 tetramer; in both systems, the tetramer remained intact throughout the trajectory with no subunit dissociation or large-scale unfolding. The lower root mean square fluctuation (RMSF) values measured for HibK386 over the equilibrated portion of the production trajectory (110–490 ns) suggest that it occupies a more rigid state than WT G6PD (Supplementary Figure 7C). Tetrameric HibK386 G6PD also had a lower solvent-accessible surface area (SASA) than WT G6PD and, consistently, formed fewer H-bonds with water molecules (3894 *±* 47 vs. 3977 *±* 45; Supplementary Figure 7D and 7E). Principal component analysis (PCA) of the WT and HibK386 trajectories with projections along PC1 and PC2 revealed that HibK386 G6PD sampled a more restricted conformational space than WT G6PD, indicating more confined dynamics (Figure 4A). Furthermore, mapping the dominant PC1 motions onto the three-dimensional structures showed that WT G6PD exhibited collective movements involving the catalytic NADP^+^ binding site and the tetramer interface, many of which were missing in HibK386 G6PD (Supplementary Figure 8A). These differences in collective movements imply that K386 2-hydroxyisobutyrylation may exert long-range effects on active-site and tetramer-interface dynamics.

**Figure 4.**
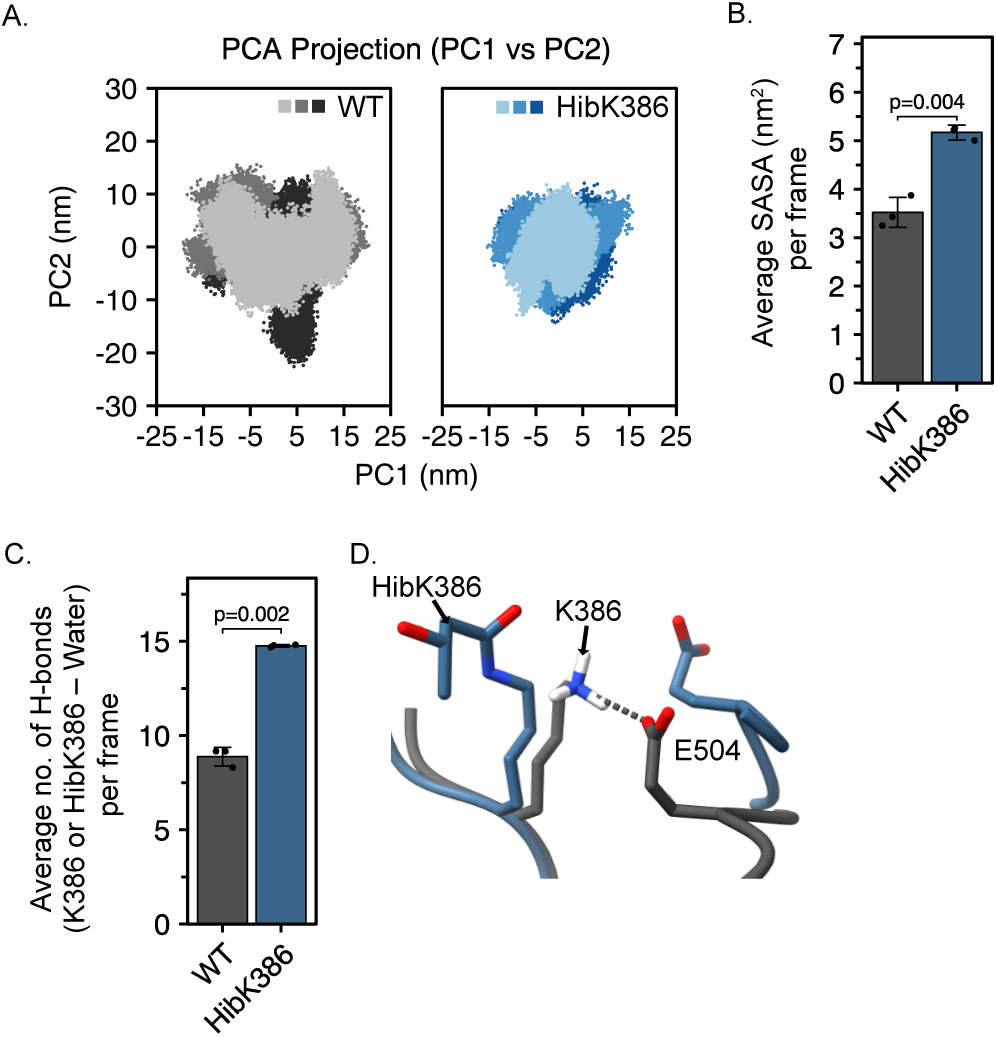
2-hydroxyisobutyrylation of K386 restricts G6PD dynamics and increases polar interactions with water molecules. **A.** Principal component analysis of the WT and HibK386 MD trajectories, projected onto the first two principal components (PC1 and PC2). Individual simulation replicas are shown in different shades of gray (WT) or blue (HibK386). **B.** Average SASA of residue 386 in WT and HibK386 G6PD, calculated in the presence of the protein matrix. Bars represent the mean *±* SD; *n* = 3 independent replicates. **C.** Average number of H-bonds formed between residue 386 and water molecules. Bars represent the mean *±* SD; *n* = 3 independent replicates. **D.** Superposition of WT (dark gray) and HibK386 (steel blue) G6PD, with K386 (or HibK386) and E504 rendered as sticks and salt bridges found only in WT G6PD marked by a dashed line. Statistical analysis was performed using a t-test (B & C).

Next, we investigated the local changes that follow the 2-hydroxyisobutyrylation of K386. At physiological pH, unmodified lysine carries a net positive charge and can form ionic interactions (salt bridges) and up to three H-bonds. 2-hydroxyisobutyrylation renders the lysine side chain neutral and adds two methyl groups, while its carbonyl and hydroxyl groups can together form up to five H-bonds, which may modify the interactions of the side chain with the solvent and with other residues (Supplementary Figure 1A). Consistent with this, the average SASA of residue 386 increased from 3.5 nm^2^ in the WT protein to 5.2 nm^2^ in HibK386 G6PD (Figure 4B and Supplementary Figure 8B). Analysis of H-bonding revealed that the average number of H-bonds formed between residue 386 and surrounding protein residues (i.e., the total number of H-bonds over the production trajectory divided by the number of frames) was lower in HibK386 compared to K386 (1.0 vs. 2.2; Supplementary Figure 8C). Conversely, the average number of H-bonds with surrounding solvent molecules increased significantly, from 8.8 for K386 to 14.7 for HibK386 (Figure 4C and Supplementary Figure 8D), demonstrating a clear preference for side chain-solvent interactions. Importantly, in WT G6PD, K386 formed a salt bridge with E504, whereas 2-hydroxyisobutyrylation abolished this strong electrostatic interaction in HibK386 G6PD (Figure 4D). These data indicate that 2-hydroxyisobutyrylation of K386 promotes interactions of residue 386 with solvent molecules at the expense of interactions with other protein side-chains, such as the stabilizing salt bridge with E504.

The G6PD tetramer is organized as a dimer of dimers, in which the assembly of the two primary dimers (chains A/B and C/D, Figure 5A) is stabilized by polar interactions across the tetramer interface, while the binding of structural NADP^+^ is essential for G6PD oligomerization and activity^20^; both oligomerization and activity were compromised in HibK386 G6PD. Structurally, K386 is located at the distal end of an extended loop, in close proximity to the structural NADP^+^ and to the dimer interface^22^ (Supplementary Figure 1B). We therefore investigated how the 2-hydroxyisobutyrylation-dependent perturbation of the local microenvironment affects the oligomeric structure of G6PD. H-bonding analysis showed that in HibK386 G6PD, the structural NADP^+^ formed more H-bonds with the protein but fewer H-bonds with surrounding water molecules (Supplementary Figure 9A and 9B). The average total number of protein-protein H-bonds was also higher in HibK386 than in WT G6PD (304 *±* 13 vs. 276 *±* 11; Figure 5B and Supplementary Figure 9C), an increase attributable mainly to intra-subunit H-bonds (Figure 5B and Supplementary Figure 9D). Analysis of inter-subunit H-bonding revealed that HibK386 G6PD formed more protein-protein H-bonds across the dimer interface but fewer H-bonds across the tetramer interface (Figures 5A and 5B, and Supplementary Figures 9E and 9F). Similarly, the average number of salt bridges between the two dimers was lower in HibK386 G6PD (9.8 *±* 1.9) than in WT G6PD (14.2 *±* 3.5), emphasizing the negative effect of K386 2-hydroxyisobutyrylation on the formation of stabilizing interactions across the tetramer interface (Figure 5C and Supplementary Figure 9G). These data indicate that K386 2-hydroxyisobutyrylation increases the number of intra-subunit and dimer-interface H-bonds, but decreases the number of H-bonds and salt bridges across the tetramer interface, potentially compromising the integrity of the G6PD tetramer.

**Figure 5.**
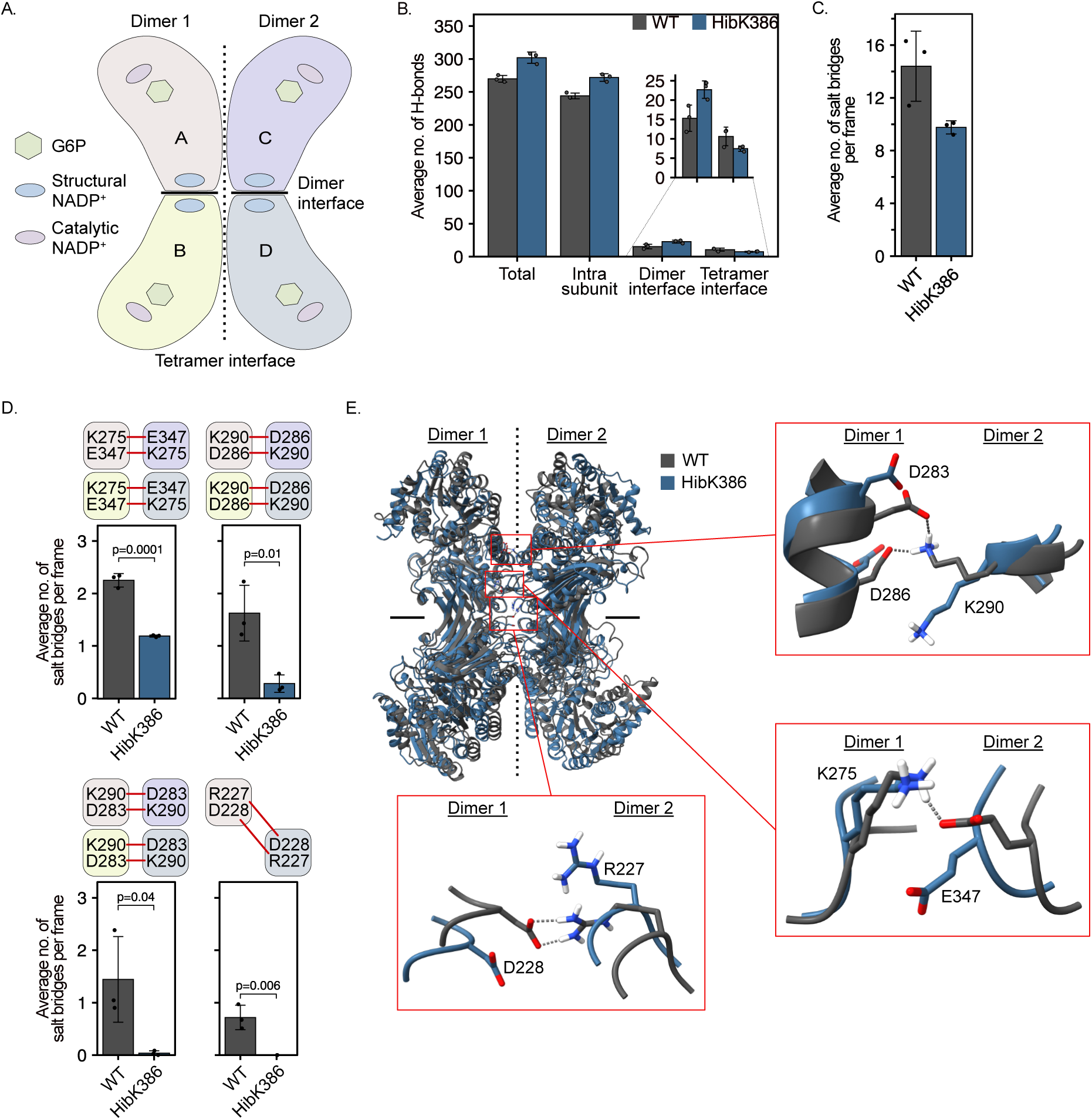
2-hydroxyisobutyrylation of G6PD at K386 weakens polar interactions across the tetramer interface. **A.** Schematic representation of the G6PD tetramer, composed of chains A and B (dimer 1) and chains C and D (dimer 2). Each protomer contains one catalytic G6P site, one catalytic NADP^+^ site, and one structural NADP^+^ binding site. The tetramer interface is marked schematically by a dotted vertical line, and the dimer interface by a solid horizontal line. **B.** Average number of protein-protein H-bonds in tetrameric G6PD. Bars represent the mean *±* SD of three independent replicates, for the total number of protein-protein H-bonds and for the individual H-bond sub-types. **C.** Average number of salt bridges formed across the tetramer interface. Bars represent the mean *±* SD; *n* = 3 independent replicates. **D.** Average number of polar interactions between selected residue pairs across the tetramer interface. Bars represent the mean *±* SD; *n* = 3 independent replicates. **E.** Superposition of WT (dark gray) and HibK386 (steel blue) G6PD, with key residue pairs at the tetramer interface rendered as sticks and polar interactions found only in WT G6PD marked by a dashed line. The focused views highlight residue pairs D283/D286–K290, K275–E347, and D228—R227. Statistical analysis was performed using a t-test (C & D).

Using the calculated principal components as collective variables, we generated a free-energy landscape to map the stable conformations sampled along the production trajectory (Supplementary Figure 9H). Detailed structural analysis of these conformations identified multiple interactions across the tetramer interface that are disrupted in HibK386 G6PD, including parallel interactions between adjacent chains (A–C and B–D) and diagonal interactions (between chains A–D and B–C). For instance, the interaction between K275 and E347 is essential for tetramer stability^22^, yet the average number of K275–E347 interactions was significantly lower in HibK386 than in WT G6PD (Figures 5D and 5E). Similarly, the parallel interactions between K290 and D286 and between K290 and D283 were disrupted in HibK386 G6PD by inward movements of the amino acid side chains. Notably, we identified an interaction between R227 and D228, formed diagonally between chains A and D, that was lost in HibK386 G6PD. Thus, MD simulations show that K386 2-hydroxyisobutyrylation results in a less dynamic structure, characterized by more polar interactions within each subunit and across the dimer interface, but by fewer stabilizing interactions across the tetramer interface. Hence, 2-hydroxyisobutyrylation of K386 close to the dimer interface results in redistribution of polar interactions at the more distant tetramer interface.

### SIRT2 is a de-2-hydroxyisobutyrylase that restores G6PD activity

Our observation that K386 2-hydroxyisobutyrylation inhibited G6PD activity only when cells were cultured in the presence of KDACi (Figure 2B) suggested that HibK386 G6PD is de-2-hydroxyisobutyrylated by endogenous deacylases. We therefore investigated the de-2-hydroxyisobutyrylation of HibK386 G6PD by NAD^+^-dependent sirtuins, as members of this family have been suggested to possess de-2-hydroxyisobutyrylase activity^7,10,11^. To this end, each of the human sirtuins SIRT1–7 was co-expressed with HibK386 G6PD in a CobB-deficient *E. coli* BL21 strain, and G6PD 2-hydroxyisobutyrylation levels were monitored by immunoblotting (Figure 6A)^42^. SIRT2 and SIRT7 efficiently catalyzed the de-2-hydroxyisobutyrylation of HibK386 G6PD in bacteria, whereas SIRT1 exhibited only modest activity. We focused on SIRT2 in light of its high de-2-hydroxyisobutyrylase activity, its recently suggested moderate activity against ACTR3^7^, and its predominantly cytoplasmic localization^43^.

**Figure 6.**
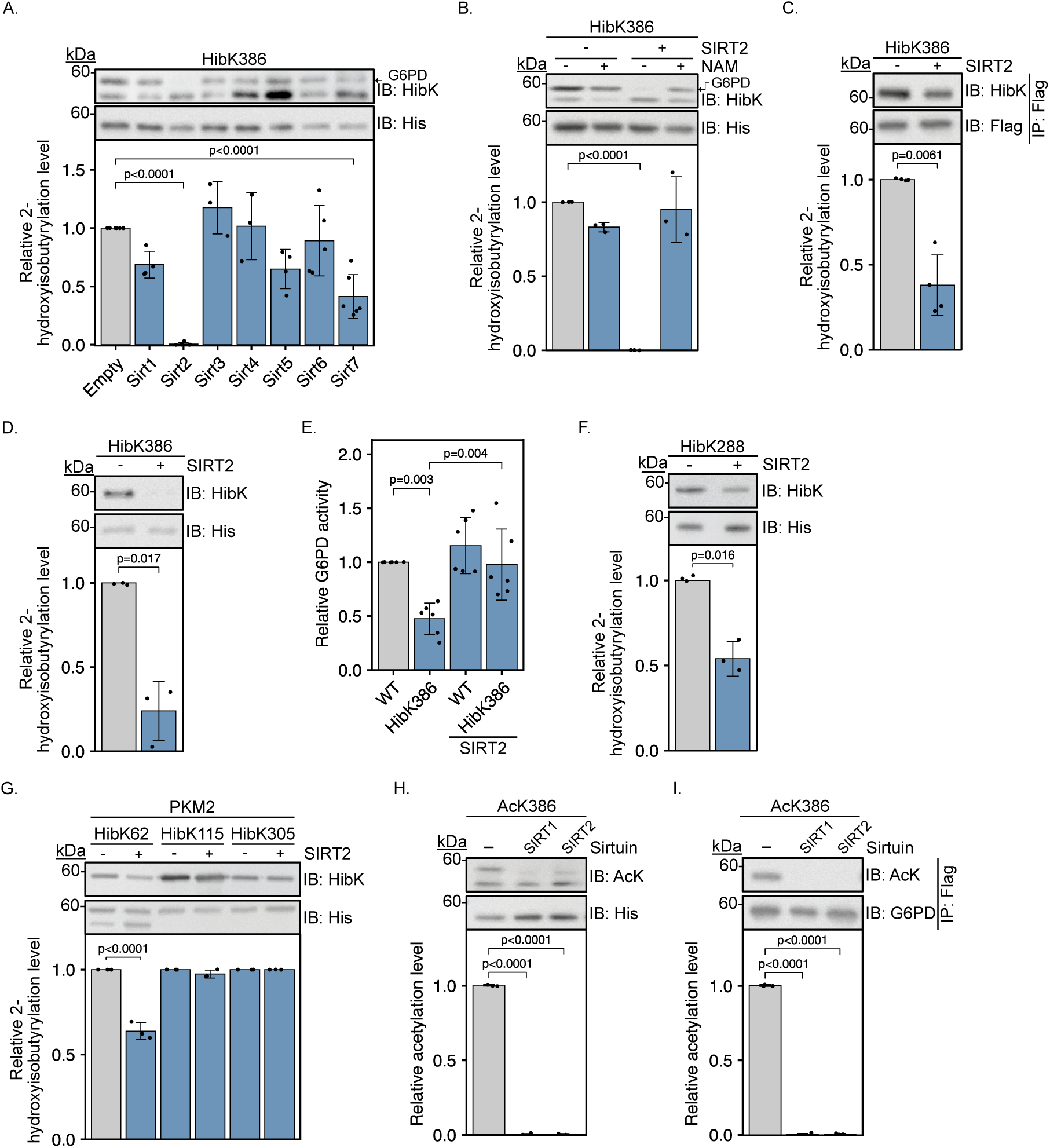
SIRT2 mediates G6PD de-2-hydroxyisobutyrylation and reactivation. **A.** de-2-hydroxyisobutyrylation of HibK386 G6PD by human sirtuins in bacteria. HibK386 G6PD was co-expressed in bacteria with SIRT1–7, and K386 2-hydroxyisobutyrylation (anti-HibK) and G6PD expression levels (anti-His) were assessed by Western blotting. Levels of K386 2-hydroxyisobutyrylation were calculated from anti-HibK immunoblot intensity relative to protein expression level, and normalized to 2-hydroxyisobutyrylation levels of HibK386 expressed in bacteria co-transformed with an empty vector. Bars represent the mean *±* SD; *n ≥* 4. **B.** SIRT2-mediated de-2-hydroxyisobutyrylation of HibK386 G6PD in bacteria. The de-2-hydroxyisobutyrylation activity of SIRT2 was assessed as described in panel A. Bacteria were cultured in the absence (-) or presence (+) of the sirtuin inhibitor NAM. Bars represent the mean *±* SD; *n* = 3. **C.** SIRT2-mediated de-2-hydroxyisobutyrylation of HibK386 G6PD in mammalian cells. Representative immunoblots show K386 2-hydroxyisobutyrylation levels (anti-HibK) and total amount of G6PD (anti-Flag), following co-expression with SIRT2 in HEK293T cells and immunoprecipitation with anti-Flag antibodies. Levels of K386 2-hydroxyisobutyrylation were calculated from anti-HibK immunoblot intensity relative to protein amount, and normalized to the 2-hydroxyisobutyrylation level of HibK386 G6PD expressed in cells co-transfected with an empty vector. Bars represent the mean *±* SD; *n* = 4. **D.** SIRT2-mediated de-2-hydroxyisobutyrylation of HibK386 G6PD in vitro. Representative immunoblots show 2-hydroxyisobutyrylation levels (anti-HibK) and the amount of purified HibK386 G6PD (anti-His) following incubation without or with purified SIRT2. Levels of K386 2-hydroxyisobutyrylation were calculated from anti-HibK immunoblot intensity relative to protein amount, and normalized to HibK386 G6PD incubated without SIRT2. Bars represent the mean *±* SD; *n* = 3. **E.** SIRT2 reactivates HibK386 G6PD. WT or HibK386 G6PD were co-expressed with SIRT2 in bacteria, as described in panel A. The activity of purified G6PD was measured under *V*_max_ conditions and normalized to the activity of WT G6PD expressed in bacteria co-transformed with an empty vector. Bars represent the mean *±* SD; *n* = 6. **F.** SIRT2-mediated de-2-hydroxyisobutyrylation of HibK288 G6PD in bacteria. The de-2-hydroxyisobutyrylation activity of SIRT2 was assessed as described in panel A. Bars represent the mean *±* SD; *n* = 3. **G.** HibK62 PKM2 is a substrate of SIRT2. PKM2 was co-expressed with SIRT2 in bacteria, and 2-hydroxyisobutyrylation levels were evaluated as described in panel A. Values are displayed relative to the 2-hydroxyisobutyrylation level of PKM2 expressed in bacteria co-transformed with an empty vector. Bars represent the mean *±* SD; *n* = 3. **H, I.** SIRT1- and SIRT2-mediated deacetylation of AcK386 G6PD in bacteria (H) and mammalian cells (I). Levels of K386 acetylation were determined using anti-AcK antibody as described in panels A and C. Bars represent the mean *±* SD; *n* = 3. Statistical analysis was performed using one-way ANOVA followed by Tukey’s post hoc test (A, B, E, G, H, and I) or t-test (C, D & F).

The de-2-hydroxyisobutyrylation of HibK386 G6PD in bacteria was SIRT2-dependent and inhibited by NAM (Figure 6B). Moreover, overexpression of SIRT2 in HEK293T cells resulted in significantly lower levels of K386 2-hydroxyisobutyrylation (Figure 6C), demonstrating that HibK386 G6PD is a substrate of SIRT2 in mammalian cells as well. In vitro de-2-hydroxyisobutyrylation assays, in which recombinant HibK386 G6PD was incubated with purified SIRT2, confirmed that SIRT2 catalyzes the de-2-hydroxyisobutyrylation of HibK386 G6PD directly (Figure 6D). G6PD K386 2-hydroxyisobutyrylation levels can therefore be regulated by SIRT2. Considering the inhibitory effect of K386 2-hydroxyisobutyrylation, we next evaluated whether SIRT2-mediated de-2-hydroxyisobutyrylation of HibK386 can reactivate the enzyme. Activity assays following co-expression in bacteria revealed that SIRT2-dependent de-2-hydroxyisobutyrylation restored G6PD activity almost to WT levels (Figure 6E). Thus, these data show that SIRT2 can recognize HibK386 G6PD, catalyze its de-2-hydroxyisobutyrylation, and restore its in vitro activity.

In addition, we measured the activity of SIRT2 against HibK288 G6PD, which we found to be inactive despite forming stable oligomers (Figure 2A–C). Immunoblot analysis following coexpression of SIRT2 with HibK288 G6PD revealed a *∼*50% reduction in K288 2-hydroxyisobutyrylation (Figure 6F), demonstrating that the de-2-hydroxyisobutyrylation activity of SIRT2 is not limited to HibK386. Moreover, to determine whether this activity extends to other proteins, we measured it against the muscle isoform of human pyruvate kinase (PKM2), site-specifically 2-hydroxyisobutyrylated at the previously identified 2-hydroxyisobutyrylation sites K62, K115, and K305^30^. Co-expression of SIRT2 with 2-hydroxyisobutyrylated PKM2 followed by immunoblot quantification revealed no activity against PKM2 HibK115 or HibK305, but an approximately 40% reduction in K62 2-hydroxyisobutyrylation (Figure 6G). Taken together, our data show that SIRT2 is a robust, yet specific, de-2-hydroxyisobutyrylase.

Having established HibK386 as a substrate of SIRT2, and given that residue K386 is subject to both acetylation and 2-hydroxyisobutyrylation, we asked whether SIRT2 can also catalyze the deacetylation of K386-acetylated G6PD. As a control, we included SIRT1, which has been reported to catalyze the deacetylation of AcK403 G6PD^29^ and the de-2-hydroxyisobutyrylation of ACTR3^7^. Deacetylation assays in bacteria showed that AcK386 G6PD is efficiently deacetylated by both SIRT1 and SIRT2 (Figure 6H), and the same was observed in mammalian cells (Figure 6I). Hence, our data suggest that SIRT2 efficiently deacylates both AcK386 and HibK386 G6PD, whereas SIRT1 is more selective, deacetylating AcK386 more efficiently than it de-2-hydroxyisobutyrylates HibK386.

Collectively, these findings establish SIRT2 as an efficient eraser of both acetyl and Hib groups at K386 that can restore HibK386 G6PD in vitro activity, and demonstrate that its de-2-hydroxyisobutyrylation activity is position-dependent across different substrates.

## Discussion

The identification of lysine 2-hydroxyisobutyrylation has expanded the ‘acylation code’ and suggested an additional potential link between cellular metabolic status and protein function. Although proteomic studies have cataloged thousands of 2-hydroxyisobutyrylation sites, the structural and functional consequences of individual modifications are mostly elusive. By genetically encoding HibK, we produced homogeneous, quantitatively modified G6PD carrying the authentic acyl group at nine defined positions and characterized the functional consequences of modification at each site. We found that 2-hydroxyisobutyrylation of K171, K205, and K360, which lie within or adjacent to the catalytic pocket, abolished activity, as did 2-hydroxyisobutyrylation of K288 at the tetramer interface. 2-hydroxyisobutyrylation of K386, positioned close to the structural NADP^+^ site and the dimer interface, reduced catalytic efficiency, destabilized the enzyme, reduced its oligomeric state, and was one of the sites at which the modification level was appreciably sensitive to endogenous KDAC activity. Comparison of K386 acetylation and 2-hydroxyisobutyrylation revealed that both modifications were destabilizing, but only 2-hydroxyisobutyrylation measurably inhibited catalysis. The two marks were also differentially recognized by their erasers: SIRT2 efficiently removed both, whereas SIRT1 was markedly more efficient against AcK386 than against HibK386, although SIRT1 can catalyze the hydrolysis of both acetyl and Hib groups. Thus, acyl identity, rather than merely the position of the acylated lysine, determines the functional outcome.

Our observations are in good agreement with the current understanding of G6PD regulation, in which oligomerization and binding of structural NADP^+^ are principal determinants of G6PD activity and stability. Stabilizing G6PD in the tetrameric state improves the activity of clinical variants, whereas loss of structural NADP^+^ binding prevents tetramerization and destabilizes the enzyme^20^. The same regulatory axis explains the pathogenicity of natural variants, whose stability and activity are tightly coupled^41^. Acetylation of K403, within the structural NADP^+^ site, abolishes activity and is reversed by SIRT2 and SIRT1^27–29^; structural and biochemical characterization of K403-acetylated G6PD suggest loss of structural NADP^+^ binding and compromised tetramer assembly^20,29^. HibK386 extends this pattern to a residue outside the cofactor pocket itself, and its functional impact is consistent with the clinical G6PD Iowa variant, in which a lysine-to-glutamate substitution at the same position impairs cofactor binding and causes severe deficiency^36^. Together, these observations support the conclusion that maintaining a positively charged lysine at position 386 is important for normal G6PD function. That said, the differential effect of K386 acetylation and 2-hydroxyisobutyrylation indicates that the chemical identity of the acetyl group, and not only the loss of positive charge, is an important factor. One may find this observation analogous to histone acylation, where different marks at a given site can have distinct functional outcomes.

The inhibition of G6PD activity by K288 2-hydroxyisobutyrylation, despite the increased proportion of higher-order oligomeric species detected by native PAGE, suggests that G6PD oligomerization is not necessarily sufficient for activity. Several studies of WT and mutated G6PD, including G6PD variants associated with clinical deficiency, have highlighted the role of structural NADP^+^ in the assembly of stable oligomers^20,23,44^. The binding of structural NADP^+^ results in conformational changes involving the C-terminus and dimer interface, which promote an oligomeric state primed for binding of G6P and catalytic NADP^+^. Wei et al.^23^ have recently suggested that oligomerization may have a role in stabilizing G6PD rather than directly affecting its catalytic activity. Thus, one explanation compatible with our data is that K288 2-hydroxyisobutyrylation drives assembly into a stable but catalytically non-productive quaternary arrangement. In HibK288 G6PD, oligomeric state and catalytic competence may therefore be separable properties. However, additional structural and biophysical studies are required to test this hypothesis.

The MD simulations suggest that K386 2-hydroxyisobutyrylation has long-range structural effects, primarily at the tetramer interface. In WT G6PD, K386 can form a salt bridge with E504, whereas 2-hydroxyisobutyrylation abolishes this electrostatic interaction and promotes H-bonding with water molecules. The redistribution of polar interactions was associated with HibK386 protomers that were more rigid overall and formed more intra-subunit and dimer-interface H-bonds, but lost polar interactions across the tetramer interface, including the K275–E347 pair required for tetramer stability^22^. In the steady-state measurements, *K*_M_ for G6P was essentially unchanged while *k*_cat_ fell by about half, arguing against a direct occlusion of the active site and favoring a model in which 2-hydroxyisobutyrylation of K386 reduces the fraction of enzyme in the active, higher-order state.

An important consideration for enzymatic or chemical acylation is stoichiometry. Site-specific acylation is typically substoichiometric, with direct measurements of human acetylation placing the median site occupancy near 0.02%^37^. If acylation drives irreversible cellular processes such as apoptosis, even low stoichiometry may exert a significant effect that accumulates over time^29^. For a monomeric enzyme, such occupancy may be functionally negligible. However, the functional effect on an oligomeric enzyme may differ because a modified subunit is incorporated into a complex whose activity is a property of the assembly. For example, in phenylalanine hydroxylase, mixed tetramers assembled from wild-type and variant subunits display activities that deviate significantly from the expected average, demonstrating that subunit properties are transmitted across the assembled oligomer^45^. If 2-hydroxyisobutyrylated G6PD subunits behave similarly, modification at a residue such as K386 or K288 could exert a disproportionate, quasi-dominant-negative effect on total cellular G6PD output. We emphasize that this is a hypothesis motivated by our data and that it must be tested using assays with mixed oligomers.

We identified SIRT2 as a KDAC capable of removing Hib from K386 in bacteria (while restoring enzymatic activity), in mammalian cells, and in vitro with purified components. This provides site-resolved evidence for de-2-hydroxyisobutyrylase activity on modified proteins, complementing recent reports of sirtuin-mediated de-2-hydroxyisobutyrylation^7,10,11^ and the CobB-dependent reaction in prokaryotes^12^. SIRT2 is a predominantly cytoplasmic sirtuin^43^, placing it in the same compartment as G6PD and fitting a broader picture in which acetylation stoichiometry and dynamics may vary among subcellular compartments according to their resident KDACs ^46,47^. The capacity of SIRT2 to remove the Hib group from HibK288 G6PD and HibK62 PKM2, in addition to HibK386 G6PD, suggests a potential role for SIRT2 that extends across enzymes of glycolysis and the PPP.

Whether G6PD 2-hydroxyisobutyrylation is installed by an acyltransferase such as p300, KAT7, KAT2A or Tip60^5,8,9,11^, or arises non-enzymatically, remains unresolved. A non-enzymatic route is plausible in principle because acyl-CoA thioesters can spontaneously acylate lysine residues at rates that scale with acyl-CoA concentration and pH^48^. However, to date, 2-hydroxyisobutyryl-CoA has not been described as an intermediate of mammalian metabolism, and the documented sources of its precursor, 2-hydroxyisobutyrate, are microbial and xenobiotic rather than endogenous^49^. Consistent with this, 2-hydroxyisobutyrate is a gut microbiome-derived metabolite that is elevated in the urine of obese individuals and tracks with particular microbial communities^50,51^. A speculative but testable hypothesis is that host-microbiome metabolic state could modulate the 2-hydroxyisobutyrylation of central metabolic enzymes, thereby affecting their activity.

In summary, we describe a workflow that combines proteomic mining, GCE, and biochemical and computational analyses to assign function to individual 2-hydroxyisobutyrylation sites, and we apply this approach to G6PD. Our results show that 2-hydroxyisobutyrylation can affect this enzyme through both its quaternary structure and activity, that the chemical identity of the acyl group at a single lysine determines both the functional outcome and the deacylase that reverses it, and that SIRT2 can selectively erase the modification. More broadly, these findings provide a template for converting the growing catalog of 2-hydroxyisobutyrylation sites into mechanistic, site-resolved biology.

### Limitations of the study

Several limitations bound these conclusions. First, the genetic encoding of HibK produces a homogeneous population of proteins that are essentially fully modified. This methodology enables direct comparison between the modified and unmodified proteins and, at the same time, reports on the maximum effect of the modification. Since 2-hydroxyisobutyrylation levels in the cell are unknown, our data establish what site-specific 2-hydroxyisobutyrylation can do, and not the extent to which it affects the cellular G6PD pool. Similarly, the activity of SIRT7 measured by deacylation assays in bacteria reports on enzymatic capacity, and may not reflect on its relative contribution to de-2-hydroxyisobutyrylation of HibK386 under physiological conditions. Second, the simulations comprise three 490 ns trajectories per tetramer, in which all four subunits are modified. While this simulation is adequate for comparing interface interaction networks, it cannot report on the dimer–tetramer equilibrium. Third, we have not measured PPP flux, NADPH/NADP^+^ ratios, or redox phenotypes in intact cells. Future studies, using different methodologies, may be able to report on the metabolic consequences of G6PD 2-hydroxyisobutyrylation in vivo.

## Methods

### Reagents

Diaphorase (#D5540), resazurin (#199303), NAD^+^ (#N7004), NADP^+^ (#N5755), G6P (#G7879), SAHA (#SML0061), NAM (#72340), and phenylmethanesulfonyl fluoride (#P7626) were obtained from Sigma-Aldrich. Sequencing grade chymotrypsin (#11418467001) and DNase I (#10104159001) were purchased from Roche Diagnostics (Mannheim, Germany). InstantBlue Coomassie Protein Stain (#ab119211) was purchased from Abcam (Cambridge, UK). Protein quantification was performed with the QPRO-BCA Kit Standard from Cyanagen (Bologna, Italy). HisPur Ni-NTA Resin (#TS-88222) was purchased from Thermo Scientific (Waltham, MA). Aprotinin (#616370), Leupeptin (#108975), Pepstatin A (#516481), and NADPH (#481973) were obtained from EMD Millipore. DNA oligomers were obtained from Sigma-Aldrich. Enzymes and buffers for molecular biology were purchased from NEB (Ipswich, MA, USA). Unless otherwise stated, general chemicals were purchased from Sigma-Aldrich (Darmstadt, Germany) and used without further purification.

### General

The *E. coli* DH10B strain was utilized for all routine cloning and plasmid amplification. Plasmid DNA was isolated and purified using NucleoSpin columns (Macherey-Nagel, Düren, Germany) in accordance with the manufacturer’s instructions. All genetic constructs were verified by Sanger sequencing. Oligonucleotide primers used for cloning and site-directed mutagenesis are provided in Supplementary Table 1. When required, lysis buffer was supplemented with a protease inhibitor cocktail consisting of 1.2 µg/mL leupeptin, 1 µM pepstatin A, 100 µM phenylmethanesulfonyl fluoride (PMSF), and 1 µg/mL aprotinin. To inhibit deacetylases, NAM and SAHA were added to buffers at final working concentrations of 20 mM and 10 µM, respectively.

### Plasmid construction

For the expression of 2-hydroxyisobutyrylated proteins in cultured mammalian cells, a single plasmid based on the pBudCE4.1 backbone (Invitrogen) was used, as previously described^29,39^. The construct contains a C-terminally Flag-tagged human *G6PD* gene and the orthogonal translation components: a U25C mutant of the amber suppressor *pylt* gene^52^, and an evolved *Methanosarcina mazei* aminoacyl tRNA synthetase (HibKRS; PylRS-C348T/Y384F) for the incorporation of HibK^34^. The vector also includes a U6 promoter-driven shRNA expression cassette (IDT; 5’-TCAGTCGGATACACACATATT-3’) for the constitutive knockdown of endogenous *G6PD*. In-frame TAG mutations were introduced into the *G6PD* sequence at specified residues via overlapping PCR (Supplementary Table 1). Plasmids for the recombinant expression of 2-hydroxyisobutyrylated G6PD in *E. coli* were generated as previously reported^29^. Briefly, the DNA fragment encoding human *G6PD* (Gift from Xiaolu Yang Addgene plasmid # 41521; http://n2t.net/addgene:41521)^53^ with a C-terminal 6*×*His tag was subcloned into a pCDF-Duet1 plasmid that contains the U25C mutant of PylT under the control of a constitutive lpp promoter^52^. In-frame TAG mutations were introduced at identified 2-hydroxyisobutyrylation sites using overlapping PCR (Supplementary Table 1). All constructs were verified by Sanger sequencing.

### Cell culture

HEK293T cells (internal stock) were maintained in DMEM (Biological Industries, Israel) supplemented with 10% heat-inactivated fetal bovine serum (FBS), 2 mM L-glutamine, 1 mM sodium pyruvate, and 100 U/mL penicillin G sodium. Cells were cultured at 37°C in a humidified incubator with 5% CO_2_. Cultures were routinely tested and confirmed negative for mycoplasma contamination.

### Overexpression of G6PD in mammalian cells

HEK293T cells were seeded in 24-well plates at a density of 3 *×* 10^5^ cells/mL. After 24 h, the culture medium was replaced with 500 µL antibiotic-free DMEM supplemented with 2 mM HibK. Where indicated, KDACi (15 mM NAM and 3 µM SAHA) were added to the medium. All media were sterilized through a 0.22 µm syringe filter and added to cells at least 30 min before transfection. Cells were transfected using polyethylenimine (PEI). After 72 h, cells were washed twice with ice-cold 1*×* phosphate-buffered saline (PBS), flash-frozen in liquid nitrogen, and stored at -20°C until further analysis.

### Native polyacrylamide gel electrophoresis (native-PAGE)

HEK293T cells transfected with plasmids encoding site-specific 2-hydroxyisobutyrylated G6PD variants were cultured in the presence of KDACi for 3 days. Cells were harvested and washed three times with ice-cold 1*×* PBS by centrifugation at 4,000*×* g for 5 min at 4°C. During the final wash, each cell suspension was divided into two equal aliquots. Cell pellets were collected by centrifugation and stored at -80°C until use. To normalize G6PD loading across samples, one aliquot was thawed on ice and lysed in 1*×* PBS supplemented with protease inhibitor cocktail and 4 µg/mL DNase I. Lysates were clarified by centrifugation at 21,000*×* g for 10 min at 4°C, and protein concentrations in the supernatants were determined using the BCA assay. Samples were denatured in 1*×* reducing Laemmli sample buffer (Bioprep, #LSB4-R) at 95°C for 5 min and resolved on 10% SDS-polyacrylamide gels. Exogenous G6PD levels were assessed by Western blotting using an anti-Flag antibody.

The second aliquot was used to assess G6PD oligomerization by native PAGE. The cells were thawed on ice and lysed in G6PD dilution buffer supplemented with protease inhibitor cocktail and 4 µg/mL DNase I. Following 25 min incubation on ice, lysates were clarified by centrifugation at 21,000*×* g for 10 min at 4°C. Based on the G6PD levels determined from the first aliquot, normalized lysate volumes were mixed with an equal volume of 2*×* native sample buffer (62.5 mM Tris-HCl [pH 6.8], 50% glycerol, 0.1% bromophenol blue) and resolved on 9% native polyacrylamide gels. Electrophoresis was performed at 4°C in native running buffer containing 20 mM Tris and 192 mM glycine. Following native PAGE, exogenous G6PD oligomerization was analyzed by Western blotting using an anti-Flag antibody. Band intensities were quantified using ImageJ^54^. The relative abundance of higher-order oligomeric species was calculated as the signal intensity of the higher oligomeric form divided by the total G6PD signal, defined as the sum of the lower and higher oligomeric forms.

### Immunoprecipitation

Total mammalian cell lysates containing exogenous Flag-tagged protein were used for Flag immuno-precipitation. Anti-Flag M2 magnetic beads (Sigma-Aldrich, #M8823) were added to clarified lysates, and samples were incubated overnight at 4°C with gentle rotation. Beads were washed three times with 1*×* PBS, and bound proteins were eluted in 1*×* non-reducing Laemmli sample buffer (Bioprep, #LSB4-NR) at 70°C for 10 min. Dithiothreitol (DTT) was then added to the eluates to a final concentration of 1 mM before immunoblotting.

### Immunoblot analysis

Protein lysates and eluates were resolved on either 10% SDS-polyacrylamide gels supplemented with 1% 2,2,2-trichloroethanol (TCE) or native polyacrylamide gels. Following electrophoresis, total protein was visualized by UV illumination of TCE-containing gels and used to assess protein loading. Proteins were then transferred to nitrocellulose membranes (Cytiva) using a semi-dry transfer system (Power Blotter, Invitrogen). Membranes were blocked for 1 h at room temperature in 5% non-fat dry milk in TBST buffer (Bio-Lab, Israel) and incubated overnight at 4°C with the indicated primary antibodies. After three washes with TBST, membranes were incubated with HRP-conjugated secondary antibodies for 1 h at room temperature. Proteins were detected using ECL reagent (Cytiva, #RPN2106) and imaged with a Fusion FX imaging system (Vilber, France).

The following primary antibodies were used for Western blotting: anti-His Tag (Applied Biological Materials, #G020; 1:4000), anti-G6PD (Abcam, #ab993; 1:5000), anti-acetyllysine (Cell Signaling Technology, #9681; 1:1000), anti-HibK (PTM BIO Inc., #PTM-801, 1:10000), anti-Flag (Sigma-Aldrich, #F1804; 1:1000). The following secondary antibodies were used for Western blotting: Goat Anti-Mouse IgG H&L (HRP) preadsorbed (Abcam, #ab7068; 1:10000), Recombinant Protein G (HRP) (Abcam, #ab7460; 1:5000), Goat Anti-Rabbit IgG H&L (HRP) (Abcam, #ab6721; 1:5000).

### Expression of recombinant G6PD in *E. coli*

Recombinant G6PD variants (WT, AcK386, and HibK386) were expressed in *E. coli* B95^55^. Cells were co-transformed with a pCDF vector encoding tRNA^Pyl^ and either WT G6PD or the K386TAG G6PD mutant and with a pBK vector encoding the evolved aminoacyl-tRNA synthetase AcKRS3 or HibKRS for site-specific incorporation of AcK or HibK, respectively^33,34^. Transformants were cultured overnight at 37°C in Luria-Bertani (LB) medium containing 50 µg/mL spectinomycin and 50 µg/mL kanamycin. The overnight cultures were used to inoculate 1 L Terrific Broth (TB) medium at an initial OD_600_ of 0.05. TB medium was supplemented with 50 µg/mL spectinomycin, 50 µg/mL kanamycin, and either 2 mM HibK or 10 mM AcK, as indicated. Cultures were grown at 37°C until the OD_600_ reached approximately 2.0. Recombinant protein expression was then induced by addition of 1 mM isopropyl β-D-thiogalactopyranoside (IPTG), and cultures were incubated for an additional 18 h at 25°C. Cells were harvested by centrifugation at 4,750*×* g and stored at -80°C until further use.

### Purification of recombinant G6PD

Frozen bacterial pellets were resuspended in Ni-NTA buffer (25 mM HEPES, pH 8.0, 150 mM NaCl, 20 mM imidazole) supplemented with protease inhibitors, 20 µg/mL DNAse I, and 20 mM NAM. Following lysis by sonication, the crude extract was clarified by centrifugation at 10,000*×* g for 45 min at 4°C. The supernatant was loaded onto a 5 mL HisTrap HP column (Cytiva, Marlborough, MA) pre-equilibrated with Ni-NTA buffer. Bound proteins were eluted using a linear gradient (0–100%) of Ni-NTA elution buffer (25 mM HEPES, pH 8.0, 150 mM NaCl, 500 mM imidazole). Fractions containing G6PD were identified by SDS-PAGE, and concentrated using a Centricon with a 30 kDa molecular weight cutoff membrane (Merck Millipore, Burlington, MA). The concentrated sample was then purified by size exclusion chromatography (SEC) using a HiLoad 26/600 Superdex 200 column (Cytiva), pre-equilibrated with SEC buffer (10 mM Tris, pH 7.5, 150 mM NaCl, 10 % glycerol). Final protein concentrations were determined by BCA assay, and the protein was stored in 30% glycerol at -80°C. The purity and identity of the recombinant enzymes were verified by SDS-PAGE followed by Coomassie staining and Western blotting, respectively.

### In bacteria de-2-hydroxyisobutyrylation assay

In-bacteria de-2-hydroxyisobutyrylation of G6PD was performed as previously described^42^. Competent CobB-knockout BL21(DE3) cells were co-transformed with plasmids for the expression of a site-specific 2-hydroxyisobutyrylated G6PD variant and the indicated deacetylase. Cells were cultured overnight at 37°C in LB medium supplemented with 50 µg/mL kanamycin, 50 µg/mL spectinomycin, and 50 µg/mL chloramphenicol. The following day, cultures were diluted to an initial OD_600_ of 0.05 in 2*×*TY medium and grown at 37°C to OD_600_ of 0.5–0.7. Cultures were then supplemented with 2 mM HibK, and protein expression was induced with 1 mM IPTG, followed by overnight incubation at 23°C. Where indicated, 20 mM NAM was added during induction to inhibit sirtuin activity. Following induction, 1 mL culture equivalents were normalized to OD_600_ of 1 using 1*×* PBS, and bacterial pellets were resuspended in 200 µL 1*×* reducing Laemmli sample buffer. Cleared lysates were obtained after incubation at 95°C for 5 min followed by centrifugation at 21,000*×* g for 10 min. G6PD 2-hydroxyisobutyrylation levels and protein expression were analyzed by Western blotting using anti-HibK and anti-His antibodies, respectively.

### G6PD activity measurement and kinetic analysis

For the measurement of G6PD activity in cell lysates, cells expressing G6PD were collected and resuspended in G6PD dilution buffer (50 mM Tris [pH 7.4], 150 mM NaCl, 0.1% NP-40) supplemented with protease inhibitor cocktail, deacetylase inhibitors, and 4 µg/mL DNase I. Lysates were clarified by centrifugation at 21,000*×* g for 10 min at 4°C. Total protein concentration in the supernatant was determined using the BCA assay. Up to 5 µg total lysate was used for enzymatic activity assays. For measurements using recombinant protein, stocks were pre-diluted to 11 nM in G6PD dilution buffer supplemented with 1 µM NADP^+^ for maintaining enzyme stability^29^.

G6PD enzymatic activity was measured using a diaphorase-resazurin coupled assay, as previously described^40^. Reactions were performed in a final volume of 100 µL containing 50 mM Tris-HCl (pH 7.4), 3.3 mM MgCl_2_, 1 U/mL diaphorase, 0.1 mM resazurin, 50 µM NADP^+^, and 1600 µM G6P. The minor contribution of NADP^+^ from the recombinant enzyme pre-dilution was included in the final NADP^+^ concentration. Reactions were initiated by the addition of cell lysate or recombinant protein (final concentration of 0.22 nM). Enzymatic activity was monitored in black 96-well plates (Greiner Bio-One) by measuring the increase in fluorescence (Ex. 530 nm, Em. 580 nm) using a microplate reader (Tecan, Männedorf, Switzerland). Initial reaction velocities were calculated from the linear portion of the resulting progress curves. To normalize activity to protein abundance in total cell lysates, reaction mixtures were collected after fluorescence measurements and analyzed by Western blotting using an anti-Flag antibody. Protein levels were quantified by densitometry using ImageJ^54^. Relative activity was calculated by normalizing the initial velocity of each reaction to the corresponding Flag-protein signal intensity and expressing the value relative to pWT. For kinetic analyses, initial velocities were measured by varying NADP^+^ concentrations from 0 to 50 µM at a fixed G6P concentration of 1600 µM, or by varying G6P concentrations from 0 to 1600 µM at a fixed NADP^+^ concentration of 50 µM. Apparent Michaelis constants (*K*_M_^app^) and maximum velocities (*V*_max_^app^) were determined by fitting initial velocity data to the Michaelis–Menten equation using R. Apparent turnover numbers (*k*_cat_^app^) were calculated from *V*_max_^app^ using a molecular weight of 59 kDa for the G6PD monomer.

### Thermal inactivation assay

WT and site-specifically 2-hydroxyisobutyrylated G6PD variants were expressed in HEK293T cells cultured in the presence of KDACi for 3 days. Cells were harvested and washed three times with ice-cold 1*×* PBS by centrifugation at 4,000*×* g for 5 min at 4°C. Cell pellets were flash-frozen in liquid nitrogen and stored at -80°C until analysis. On the day of the assay, pellets were thawed on ice and resuspended in 1*×* PBS supplemented with protease inhibitor cocktail and 4 µg/mL DNase I. No additional NADP^+^ was added to the buffer. After 25 min incubation, lysates were clarified by centrifugation at 21,000*×* g for 10 min at 4°C, and total protein concentration was determined using the BCA assay. Clarified lysates were aliquoted into PCR tubes and exposed to the indicated temperatures for 3 min using a thermal cycler. Samples were equilibrated at room temperature for 3 min, after which residual G6PD activity was measured. Thermal inactivation curves were generated by plotting initial reaction velocities as a function of temperature. The thermal stability of purified recombinant G6PD variants was measured in a similar way. Protein stocks were diluted to a working concentration of 9.8 nM in PBS supplemented with 1 µM NADP^+^. Samples were then subjected to the same thermal inactivation procedure, and residual activity was measured as described above.

### Chymotrypsin digestion

To measure the proteolytic stability of G6PD, 12.20 µg recombinant G6PD was incubated with 2.8 µg chymotrypsin in 700 µL chymotrypsin digestion buffer (100 mM Tris-HCl [pH 8.0], 10 mM CaCl_2_) at room temperature. At the indicated time points, reaction aliquots were withdrawn and immediately quenched by boiling in 1*×* reducing Laemmli sample buffer to terminate chymotrypsin activity. Remaining full-length G6PD was analyzed as a function of time by SDS-PAGE followed by Coomassie staining.

### LC-MS/MS

For mass spectrometric analysis, purified G6PD was resolved by 10% Tris-glycine SDS-PAGE and visualized with 0.1% (w/v) Coomassie Brilliant Blue R250. The protein band corresponding to G6PD was excised and subjected to in-gel reduction with 3 mM DTT at 60°C for 30 min, followed by alkylation with 10 mM iodoacetamide in 100 mM ammonium bicarbonate at room temperature in the dark for 30 min. In-gel digestion was performed with modified trypsin (Promega) at a 1:10 enzyme-to-substrate ratio in 10 mM ammonium bicarbonate containing 10% (v/v) acetonitrile at 37*^◦^*C for 16 h. The resulting peptides were extracted from the gel matrix and desalted using C18 StageTips prior to liquid chromatography-tandem mass spectrometry (LC-MS/MS). Peptides were separated by reverse-phase nano-flow chromatography using 0.075 *×* 300-mm fused silica capillaries (J&W) packed with Reprosil-Pur C18-AQ material (Dr. Maisch GmbH). Chromatography was performed at a flow rate of 0.15 µL/min using a linear gradient of Buffer A (0.1% [v/v] trifluoroacetic acid in water) and Buffer B (0.1% [v/v] trifluoroacetic acid in acetonitrile). The elution profile consisted of 5%–28% B over 30 min, 28%–95% B over 15 min, and a 15 min wash at 95% B. Mass spectra were acquired in positive ion mode using a Q Exactive plus mass spectrometer (Thermo Fisher Scientific). The instrument was operated in data-dependent acquisition mode. Full MS scans were followed by high-energy collision dissociation of the ten most abundant precursor ions. MS/MS raw files were processed using Proteome Discoverer 2.4 (Thermo Fisher Scientific) utilizing the Sequest HT search engine. Data were searched against the Human UniProt database supplemented with the sequence of the recombinant His-tagged G6PD construct. Search parameters were defined with a precursor mass tolerance of 10 ppm and a fragment ion tolerance of 0.05 Da. Static modifications included carbamidomethylation of cysteine residues. Variable modifications allowed were oxidation (M), phosphorylation (S, T, Y), acetylation (K), ubiquitination/di-glycine (K), methylation (K), and 2-hydroxyisobutyrylation (K). A maximum of two missed tryptic cleavages was permitted, and the minimum peptide length was set to six amino acids. False discovery rates (FDR) for both peptides and proteins were maintained at *<*1% using a target-decoy strategy. For semi-quantitative analysis, peptide abundance was determined by calculating the area under the curve of extracted ion chromatograms. Protein abundance was estimated by averaging the peak areas of the three most intense unique peptides. Mass spectrometry was performed at the Ilse Katz Institute for Nano-Science and Technology, Ben-Gurion University of the Negev.

### Multiple sequence and structural alignments

To identify evolutionarily conserved 2-hydroxyisobutyrylation sites in G6PD, we integrated published proteomics data on 2-hydroxyisobutyrylation across different organisms with multiple sequence alignment and superposition of protein structures. For each organism, MS data were extracted from the following sources, using the indicated UniProt accession number (in parentheses): *Homo sapiens* (P11413)^8,30,31^, *Mus musculus* (Q00612)^4^, *Oryza sativa* (Q7EYK9)^3^, *Triticum aestivum* (A0A3B6QEF1)^56^, *Physcomitrium patens* (A9SA38)^57^, *Candida albicans* (A0A1D8PEG2)^58^, *Fusarium oxysporum* (A0A0J9U458)^59^, *Aspergillus niger* (A0A505I6K0)^60^, *Aspergillus flavus* (B8N022)^61^, *Frankliniella occidentalis* (A0A6J1RR48)^62^, *Botryotinia fuckeliana* (A0A384J7Q3)**^?^**, *Ustilaginoidea virens* (A0A063C5C1)^63^, *Giardia lamblia* (A8BWM5)^64^, *Trypanosoma brucei* (Q38BZ6)^65^, and *Proteus mirabilis* (A0A2X2DKS9)^2^. Multiple sequence alignment was performed using Clustal Omega (version 1.2.2)^66^ with default parameters to assess the conservation of modified lysine residues.

Three-dimensional (3D) structures of G6PD orthologs were either retrieved from the Protein Data Bank (PDB) or modeled using AlphaFold (version 2.0)^67^, with prediction reliability confirmed by mean pLDDT scores > 90 across the aligned sequences. For structural conservation analysis, 3D structures were superimposed, visualized, and rendered using PyMOL (version 2.5.4, Schrödinger, LLC)^68^.

### Molecular dynamics

Molecular dynamics simulations were performed using GROMACS 2025.2^69,70^. The G6PD monomer structure was obtained from the PDB (ID: 6E08)^40^, and the tetramer was generated in ChimeraX 1.10.1^71^. Structural NADP^+^ was incorporated via superposition with PDB ID: 7SNI^23^, and 2-hydroxyisobutyrylation was introduced using the Vienna-PTM web server^72^. The CHARMM-36M force field was applied using CHARMM-GUI^73,74^, and systems were solvated with TIP3P water and neutralized with 0.15 M NaCl, resulting in systems of approximately 293,240 atoms in a 105*×*105*×*105 Å cubic box. Systems were minimized using the steepest-descent method, followed by equilibration with the v-rescale thermostat at 303.15 K. Three independent 490 ns production runs (1.47 µs total) were performed for each variant, using a 2.0 fs time step and a seed generator. Long-range electrostatics were calculated using PME under PBC, with a 12 Å cut-off and a 10 Å force switch. Convergence was assessed using GROMACS tools to calculate the mean Cα RMSD, RMSF, SASA, Rg, H-bonds, and PCA. Hydrogen bonds were calculated using standard GROMACS criteria (donor–acceptor distance *≤* 0.35 nm and hydrogen–donor–acceptor angle *≤* 30*^◦^*). Analyses focused on residues critical for oligomerization identified by PDBePISA^75^ and FACE2FACE^76^. Custom scripts built based on MDanalysis software were used for salt-bridge time series and free energy landscape analysis^77,78^. Salt bridges were defined as interactions between basic side-chain nitrogen atoms (Arg/Lys) and acidic side-chain oxygen atoms (Asp/Glu) within a cutoff distance of *≤* 0.4 nm.

### Statistics and reproducibility

The experiments were not randomized, and the investigators were not blinded to allocation during experiments and outcome assessment. Statistical analyses were performed in R. Multiple comparisons were made using one-way ANOVA followed by Tukey’s post hoc test; pairwise comparisons were made using a two-tailed Student’s t-test, as indicated in the corresponding figure legends. *p <* 0.05 was considered statistically significant. The number of independent biological replicates for each experiment is given in the corresponding figure legend.

## Acknowledgements

This work was supported by the European Research Council (ERC) under the European Union Horizon 2020 research and innovation programme [grant number 678461 to E.A.]; the Israel Science Foundation [grants number 1769/20 and 2331/25 to E.A.]; and the Israeli Ministry of Innovation, Science, and Technology [grant number 5936 to E.A.]

## Author contributions

Conceptualization: S.P., F.W., and E.A. Investigation: S.P., O.E.H., and F.W. Formal analysis: S.P., F.W., and E.A. Visualization: S.P. Funding acquisition: E.A. Supervision: F.W., I.K., and E.A. Writing: S.P., F.W., and E.A. All authors commented on the manuscript.

## Competing interests

The authors declare no competing interests.

## Notes

### Competing Interest Statement

The authors have declared no competing interest.

