## Supplementary Information for "One lysine, two acyl marks: 2-hydroxyisobutyrylation and acetylation of G6PD differ in outcome and regulation"

### Supplementary Figures

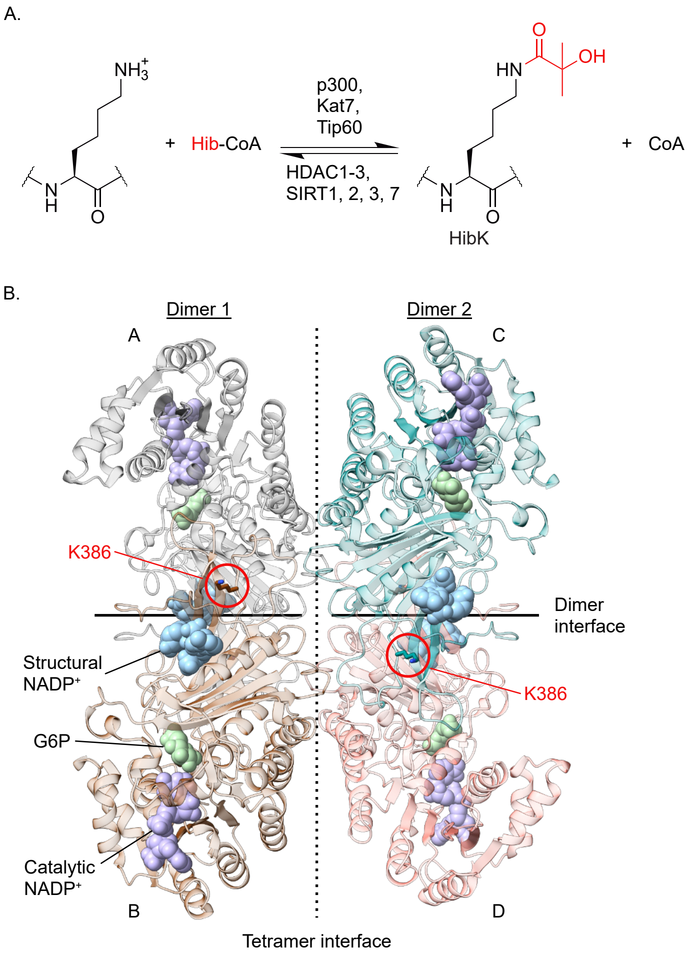


**Supplementary Fig. 1** Lysine 2-hydroxyisobutyrylation and the structure of human G6PD. **A.** Post-translational 2-hydroxyisobutyrylation of lysine residues. The acylation reaction is catalyzed by lysine acyltransferases, and the deacylation reaction by lysine deacylases. **B.** Crystal structure of inactive D200N human G6PD tetramer, showing structural $NADP^+$, the substrate G6P, and the $NADP^+$ cofactor (PDB ID: 7SNI)[1]. The tetramer and dimer interfaces are schematically indicated by dashed and continuous lines, respectively. The position of K386, close to the structural $NADP^+$ and the dimer interface, is highlighted by a red circle.

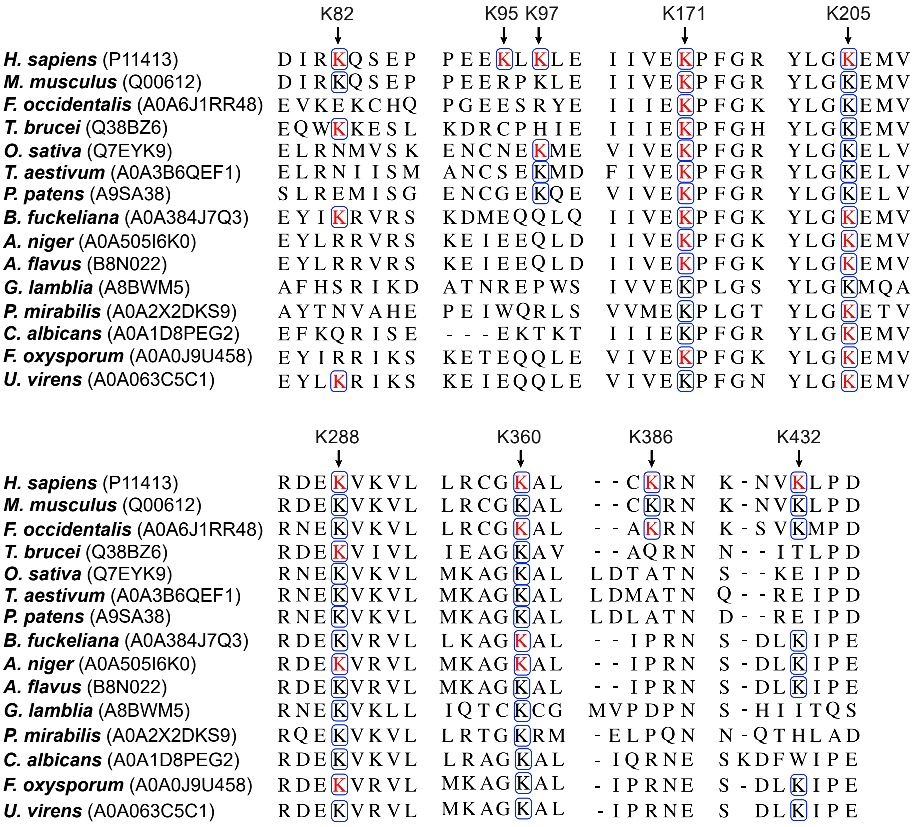


**Supplementary Fig. 2** Evolutionary conservation of G6PD 2-hydroxyisobutyrylation sites. Multiple sequence alignment of G6PD orthologs from the indicated organisms, with lysine residues identified as 2-hydroxyisobutyrylated by mass spectrometry shown in red, and evolutionarily conserved lysine residues highlighted by a blue frame. Accession numbers are given in parentheses, and residue positions are numbered according to the human G6PD sequence.

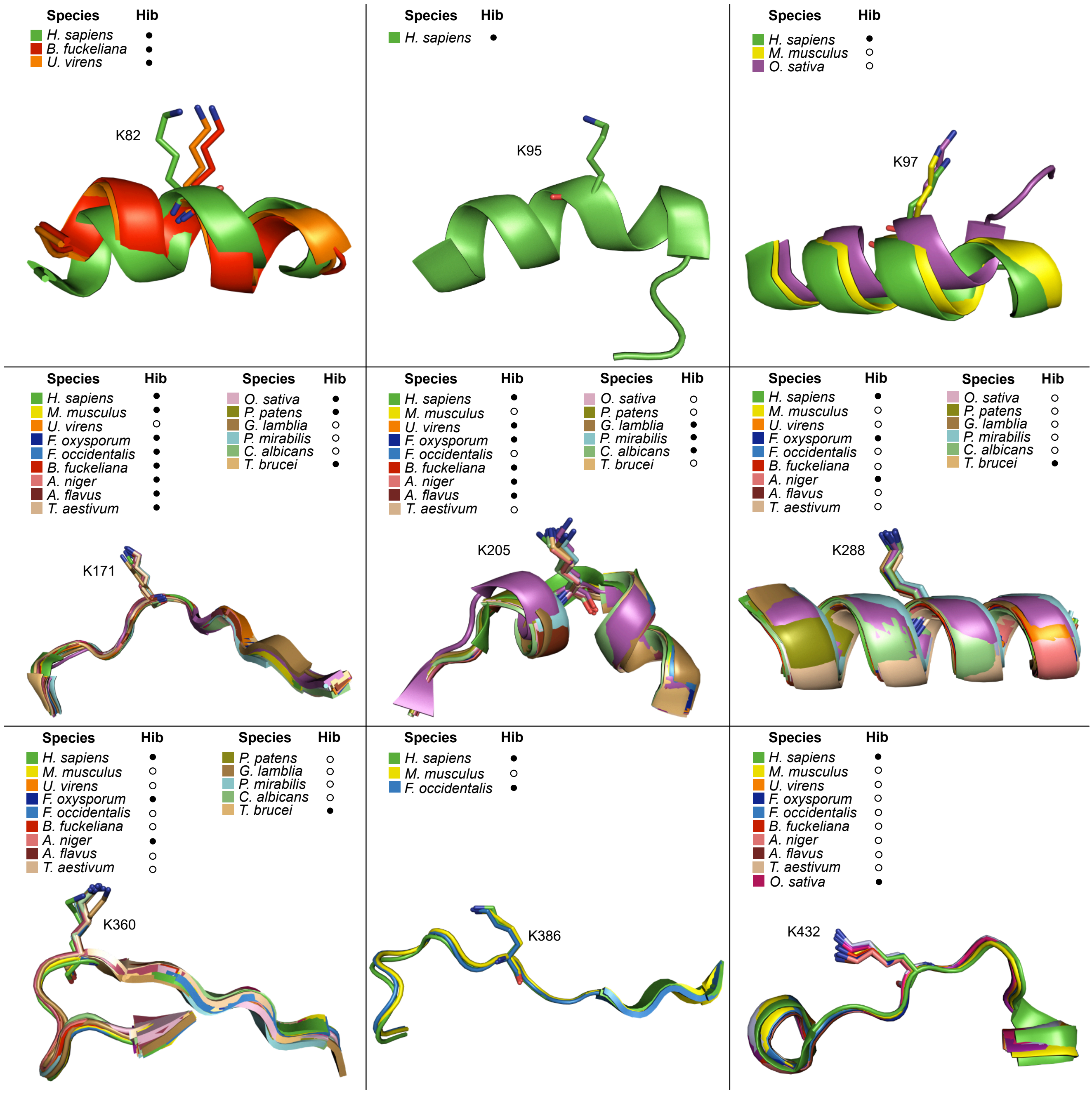


**Supplementary Fig. 3** Structural conservation of G6PD 2-hydroxyisobutyrylation sites. Superimposition of three-dimensional structures of G6PD orthologs, focused on the lysine residues investigated as potential 2-hydroxyisobutyrylation sites in human G6PD. Mass spectrometry-based 2-hydroxyisobutyrylation data were compiled across 15 organisms (UniProt accession numbers and corresponding citations are detailed in the Materials and Methods section). Dots (•) mark lysine residues identified as 2-hydroxyisobutyrylation sites in mass spectrometry studies. All structures, except for human G6PD (PDB ID: 6E08[2]), were predicted by AlphaFold 2.0 (mean pLDDT > 90 for the aligned sequence regions)[3].

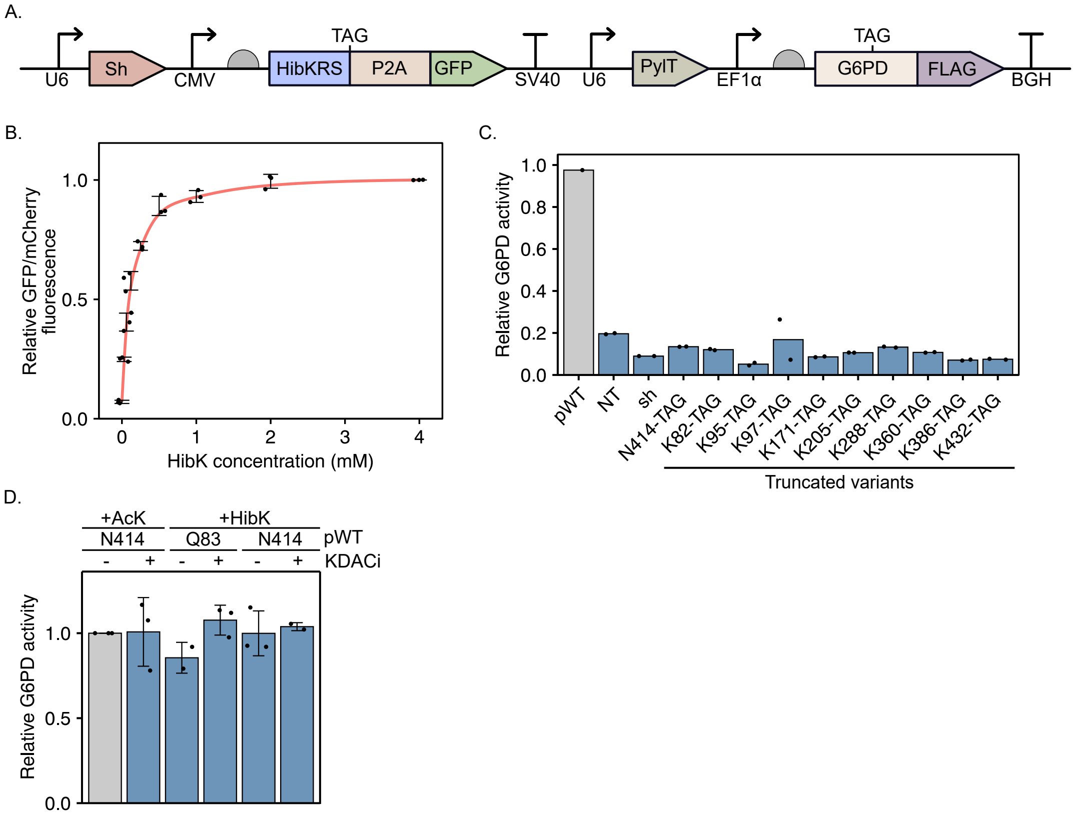


**Supplementary Fig. 4** Expression of site-specifically 2-hydroxyisobutyrylated G6PD in cultured mammalian cells. **A.** Single-plasmid system for the expression of site-specifically 2-hydroxyisobutyrylated G6PD in cultured mammalian cells. The vector carries an orthogonal HibK tRNA synthetase/pyrrolysine tRNA pair (HibKRS/$tRNA^{Pyl}_{CUA}$) for site-specific incorporation of HibK in response to an in-frame UAG stop codon, an shRNA against endogenous *G6PD*, and a C-terminally Flag-tagged, sh-resistant *G6PD* gene. **B.** Dose-dependent incorporation of HibK. HEK293T cells expressing an mCherry-TAG-GFP reporter were treated with the indicated concentrations of HibK. GFP/mCherry fluorescence ratios are presented as a function of HibK concentration, relative to the measurement at 4 mM HibK. The red line serves as a guide to the eye. Data are the mean ± SD; $n = 3$ biological replicates. **C.** Enzymatic activity of pWT and C-truncated G6PD relative to background. Truncated G6PD variants were expressed in HEK293T cells cultured without HibK, and enzymatic activity was measured in cleared cell lysates under $V_{max}$ conditions. Non-transfected (NT) and G6PD knockdown (sh) cells were included as controls. Activity is presented relative to that measured in lysates of cells expressing full-length pWT G6PD and cultured with HibK. Bars represent the mean; $n = 2$. **D.** Functional validation of HibK414 G6PD as a non-2-hydroxyisobutyrylated control. Bars represent the $V_{max}$ of exogenous G6PD measured in cleared lysates of cells cultured without or with KDACi, normalized to immunoblot intensities of G6PD expression and displayed relative to the published control (AcK414)[4]. Bars represent the mean ± SD; $n = 3$. Statistical analysis was performed using one-way ANOVA followed by Tukey's post hoc test.

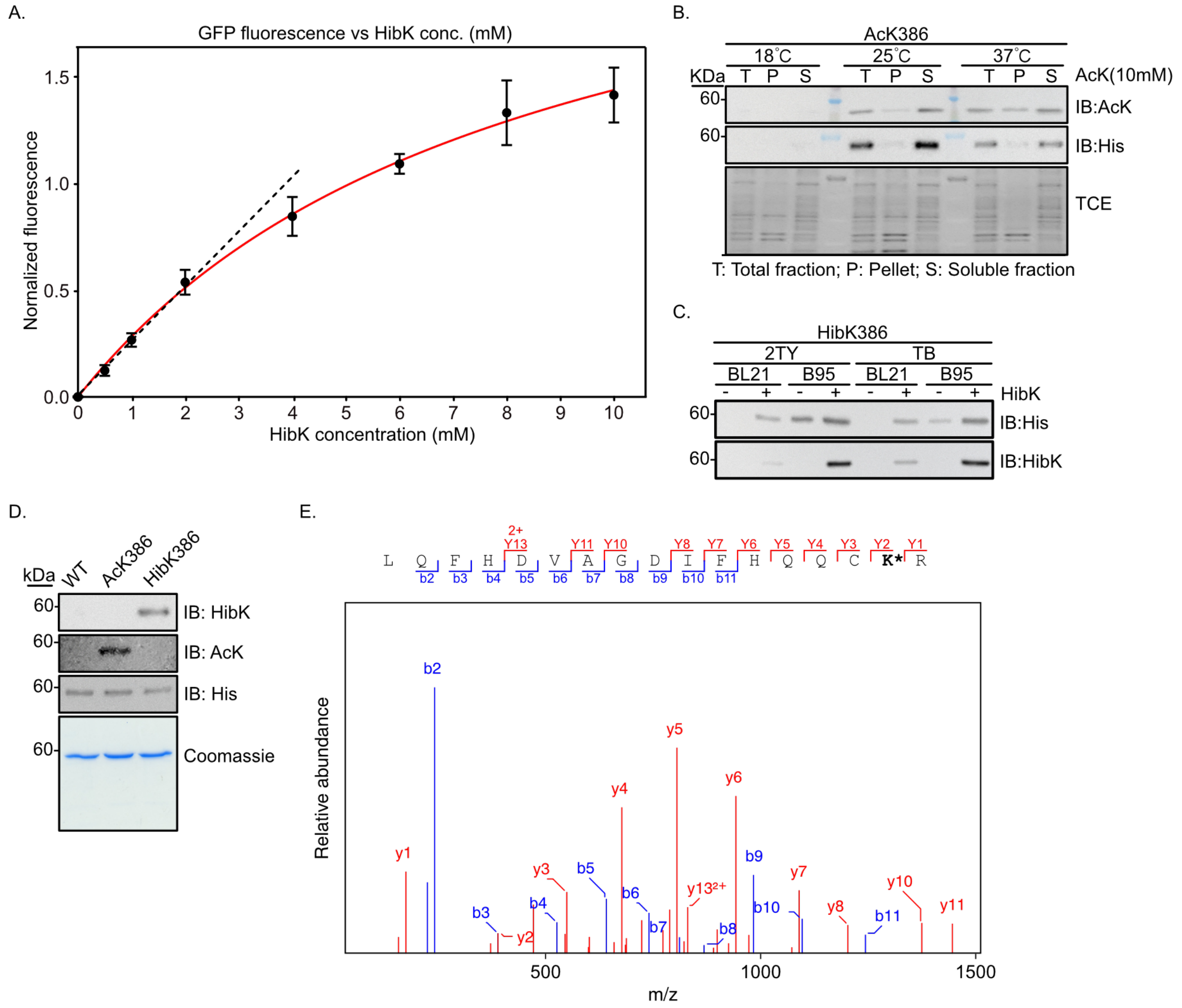


**Supplementary Fig. 5** Optimization of expression conditions in *E. coli*, protein purification, and Michaelis–Menten kinetics. **A.** HibK dose-dependent incorporation efficiency during bacterial expression. Quantification of superfolder GFP (sfGFP)-150TAG reporter fluorescence, normalized to optical density (OD), in response to increasing concentrations of HibK in the culture medium. Protein expression level increased almost linearly with HibK concentration in culture media, up to ∼2 mM. Lines serve as a guide to the eye. **B.** Evaluation of the induction temperature for G6PD expression. Immunoblotting with anti-His and anti-AcK antibodies identified 25°C as the optimal temperature for maximal soluble protein yield. **C.** Comparison of G6PD expression levels in *E. coli* BL21 and B95 strains. Expression levels were confirmed by immunoblotting with anti-His antibodies, and HibK incorporation with anti-HibK antibodies. **D.** Validation of purified WT, AcK386, and HibK386 G6PD. Site-specific incorporation and modification specificity were confirmed by immunoblotting with anti-AcK and anti-HibK antibodies. Protein purity and identity were verified by Coomassie-stained SDS-PAGE and by anti-His immunoblotting, respectively. **E.** LC-MS/MS analysis of bacterially expressed and purified 2-hydroxyisobutyrylated G6PD. The sequence and observed mass of the tryptic peptide shown confirm site-specific incorporation of HibK (indicated by K*) at the designated position. The modified peptide sequence and the measured mass of the precursor ion are presented.

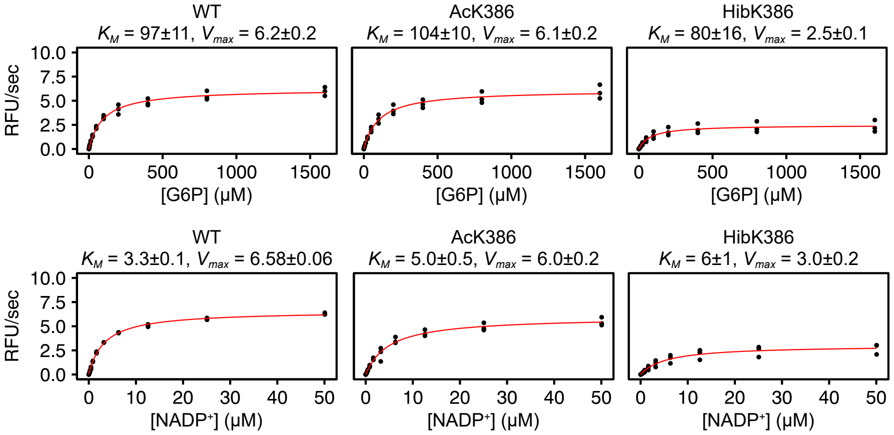


**Supplementary Fig. 6** Michaelis–Menten analysis of recombinant G6PD variants. Steady-state kinetic analysis of G6PD activity as a function of G6P (top) or $NADP^+$ (bottom) concentration. Initial rates are plotted against substrate concentration and fitted to the Michaelis–Menten equation (red line), from which $V_{max}^{app}$ and $K_M^{app}$ values were extracted. Values are reported as the mean ± fitting error; $n = 3$ independent measurements.

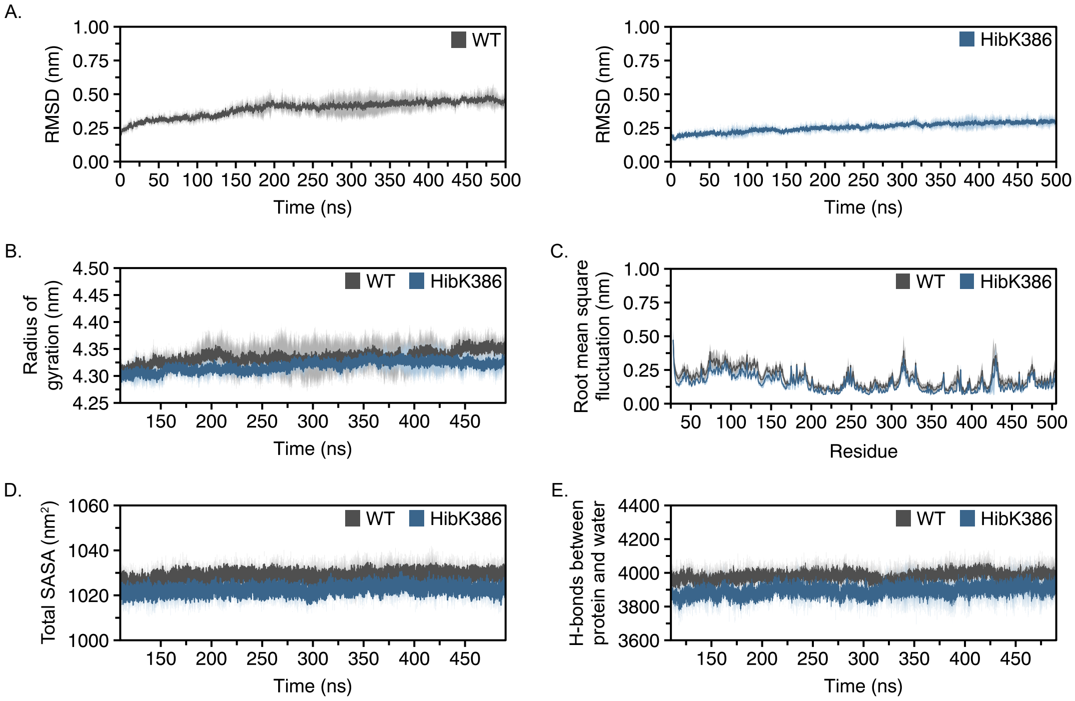


**Supplementary Fig. 7** Global structural and dynamic properties of WT and HibK386 G6PD tetramers along the MD trajectories. **A.** C$\alpha$ root-mean-square deviation (RMSD) of the WT and HibK386 G6PD tetramers relative to the initial coordinates. On the basis of the RMSD, the 7 highly flexible and unstructured C-terminal residues were truncated. **B.** Radius of gyration (Rg) of the WT and HibK386 G6PD tetramers along the trajectory. **C.** Root-mean-square fluctuation (RMSF) per residue (excluding hydrogen atoms) relative to the initial coordinates. **D.** Total solvent-accessible surface area (SASA) of the WT and HibK386 G6PD tetramers. **E.** Number of H-bonds formed between amino acid side chains and water molecules in the WT and HibK386 G6PD tetramers. Data in all the panels are presented as a function of simulation time, with solid lines representing the average across three independent simulations in dark gray (WT) and steel blue (HibK386); light gray and light steel blue shaded regions represent the standard error.

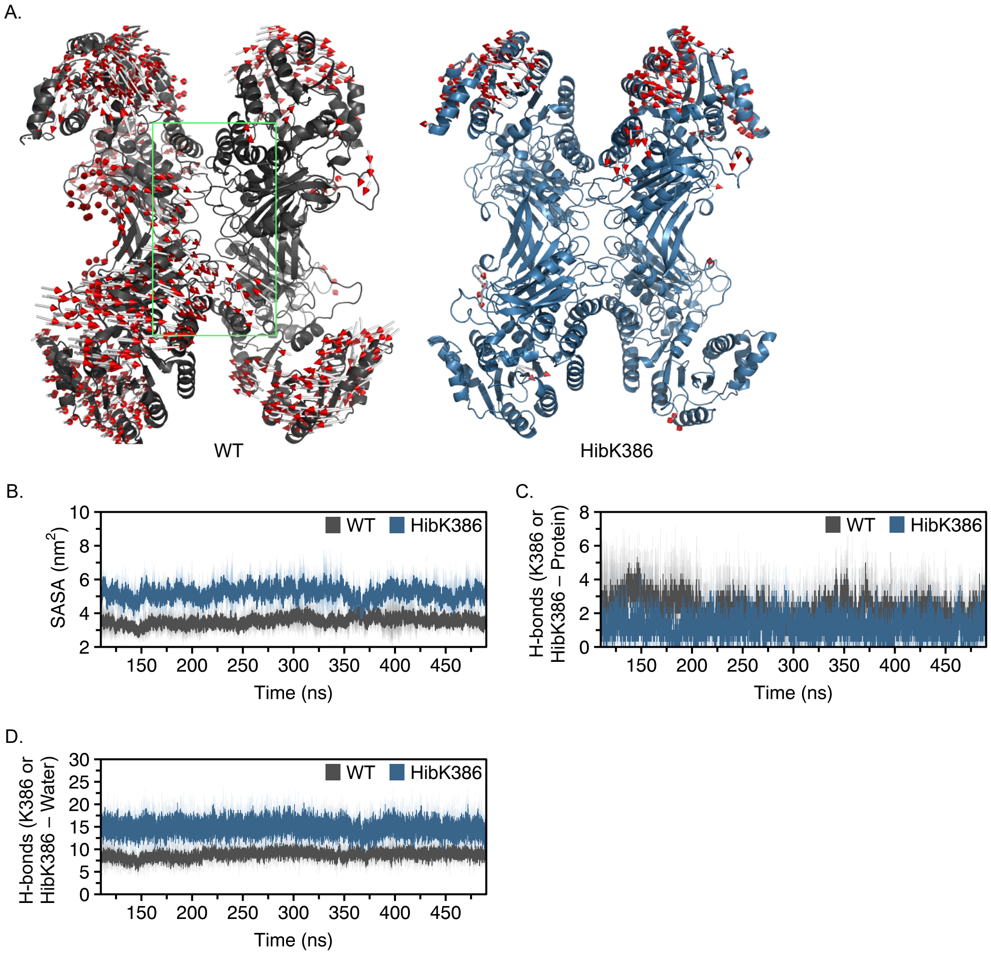


**Supplementary Fig. 8** Collective motions and the local environment of residue 386 in WT and HibK386 G6PD. **A.** Mapping of the dominant PC1 motions onto the three-dimensional structures of WT and HibK386 G6PD. Red arrows indicate the direction of the dominant motions, and the tetramer interface is outlined by a green box. Distinct motions at the tetramer interface are apparent in WT but not in HibK386 G6PD. **B.** Time-averaged SASA of residue 386 calculated in the presence of the full protein matrix (G6PD tetramer). **C.** Side chain H-bonds between residue 386 and the rest of the protein (G6PD tetramer). **D.** Side chain H-bonds between residue 386 and water in the G6PD tetramer. Data in panels B–D are presented as a function of simulation time, with solid lines representing the average across three independent simulations in dark gray (WT) and steel blue (HibK386); light gray and light steel blue shaded regions represent the standard error.

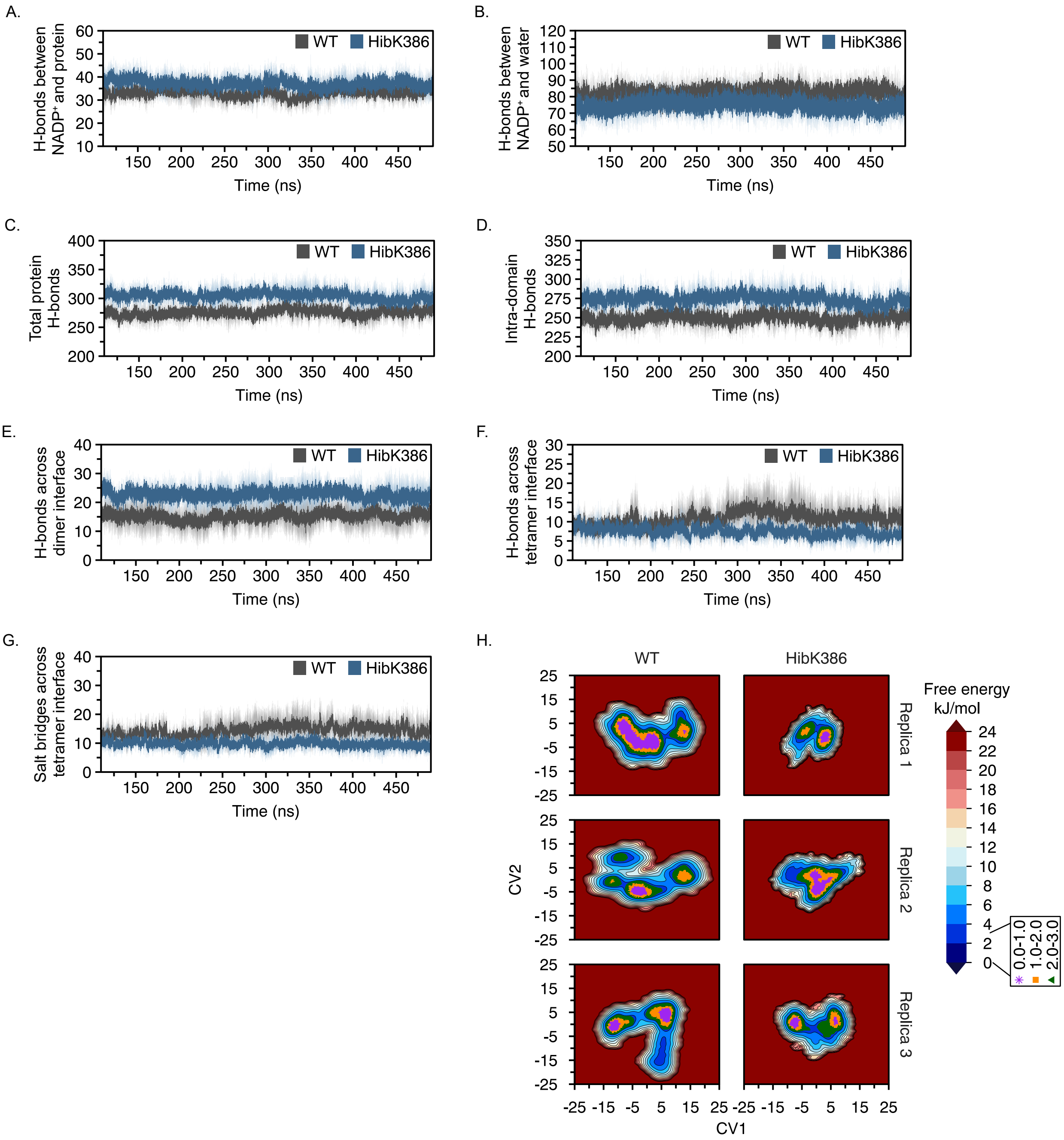


**Supplementary Fig. 9** Polar interactions within and between G6PD subunits, and free-energy landscape analysis. **A.** Number of H-bonds formed between structural $NADP^+$ and amino-acid side chains in the G6PD tetramer. **B.** Number of H-bonds formed between structural $NADP^+$ and water molecules in the G6PD tetramer. **C.** Total number of H-bonds formed between amino-acid side chains in the G6PD tetramer. **D.** Intra-subunit side chain H-bonds in the G6PD tetramer, excluding interface interactions. **E.** Amino-acid side chain H-bonds across the dimer interface in the G6PD tetramer. **F.** Amino-acid side chain H-bonds across the tetramer interface in the G6PD tetramer. **G.** Salt bridges formed between amino acid side chains across the tetramer interface in the G6PD tetramer. **H.** Free-energy landscape analysis. PC1 and PC2 from the PCA were used as collective variables (CVs) to characterize conformational sampling along the trajectory. Markers on the free-energy map highlight the principal low-energy basins: global minimum (0.0–1.0 kJ/mol, pink plus), secondary minimum (1.0–2.0 kJ/mol, yellow square), and tertiary minimum (2.0–3.0 kJ/mol, green triangle). Data in panels A–G are presented as a function of simulation time, with solid lines representing the average across three independent simulations in dark gray (WT) and steel blue (HibK386); light gray and light steel blue shaded regions represent the standard error.

### Supplementary Tables

**Supplementary Table 1** Primers used in this work.

| Oligo | Sequence |
|---|---|
| pBud/NotI_Forward | ATAAGAATGCGGCCGCATGGCAGAGCAGGTGGCCC |
| pBud/KpnI_Reverse | CGGGGTACCGAGCTTGTGGGGGTTCACCCAC |
| pBud/N414TAG_Forward | AAGCCGGGCATGTTCTTCTAGCCCGAGGAGTCGGAGCTG |
| pBud/N414TAG_Reverse | CTCCGACTCCTCGGGCTAGAAGAACATGCCCGGCTTCTTG |
| Q83TAG_Forward | GTGGCTGACATCCGCAAATAGAGTGAGCCCTTCTTCAAG |
| Q83TAG_Reverse | TGAAGAAGGGCTCACTCTATTTGCGGATGTCAGCCACTG |
| K82TAG_Forward | TGACATCCGCTAGCAGAGTGAGCCCTTC |
| K82TAG_Reverse | GGGCTCACTCTGCTAGCGGATGTCAGC |
| K95TAG_Forward | AAGGCCACCCCAGAGGAGTAGCTCAAGCTGGAGGACTTC |
| K95TAG_Reverse | GTCCTCCAGCTTGAGCTACTCCTCTGGGGTGGCCTTGAAG |
| K97TAG_Forward | ACCCCAGAGGAGAAGCTCTAGCTGGAGGACTTCTTTGCC |
| K97TAG_Reverse | CAAAGAAGTCCTCCAGCTAGAGCTTCTCCTCTGGGGTGG |
| K171TAG_Forward | ACCGCATCATCGTGGAGTAGCCCTTCGGGAGGGACC |
| K171TAG_Reverse | TCCCTCCCGAAGGGCTACTCCACGATGATGCGGTTC |
| K205TAG_Forward | CCACTACCTGGGCTAGGAGATGGTGCAGAACC |
| K205TAG_Reverse | TTCTGCACCATCTCCTAGCCCAGGTAGTGGTC |
| K288TAG_Forward | GACGTCCGTGATGAGTAGGTCAAGGTGTTGAAATGC |
| K288TAG_Reverse | ATTTCAACACCTTGACCTACTCATCACGGACGTCATC |
| K360TAG_Forward | CGCTGCGGCTAGGCCCTGAACGA |
| K360TAG_Reverse | TTCAGGGCCTAGCCGCAGCGCA |
| K386TAG_Forward | TCTTCCACCAGCAGTGCTAGCGCAACGAGCTGGTGATC |
| K386TAG_Reverse | TCACCAGCTCGTTGCGCTAGCACTGCTGGTGGAAGATG |
| K432TAG_Forward | AACAGATACAAGAACGTGTAGCTCCCTGACGCCTACGAG |
| K432TAG_Reverse | GTAGGCGTCAGGGAGCTACACGTTCTTGTATCTGTTGCC |
| pCDF/NdeI_forward | GGAATTCCATATGGCAGAGCAGGTGGCCCTGAGC |
| pCDF/His_XhoI_Reverse | CCGCTCGAGTTAGTGGTGATGATGGTGATGGAGCTTGTG GGGGTTC ACCCAC |
| pCDF/N414TAG_Forward | CGGGCATGTTCTTCTAGCCCGAGGAGTC |
| pCDF/N414TAG_Reverse | GCTCCGACTCCTCGGGCTAGAAGAACATGC |

**Supplementary Table 2** Kinetic parameters of G6PD variants. Apparent Michaelis constants ($K_M^{app}$), maximal velocities ($V_{max}^{app}$), turnover numbers ($k_{cat}^{app}$), and catalytic efficiencies ($k_{cat}^{app}/K_M^{app}$) were determined for G6P and NADP$^+$ by fitting to the Michaelis–Menten equation. Data are the mean ± SD; $n$ = 3 independent measurements.

| Variant | G6P | | | |
|---|---|---|---|---|
| | $V_{max}^{app}$ (µM·s$^{-1}$) | $K_M^{app}$ (µM) | $k_{cat}^{app}$ (s$^{-1}$) | $k_{cat}^{app}/K_M^{app}$ (µM$^{-1}$·s$^{-1}$) |
| WT | 0.023±0.002 | 98±17 | 105±8 | 1.1±0.2 |
| AcK386 | 0.023±0.002 | 104±2 | 96±11 | 0.9±0.1 |
| HibK386 | 0.009±0.002 | 80±7 | 42±11 | 0.5±0.1 |

| Variants | NADP$^+$ | | | |
|---|---|---|---|---|
| | $V_{max}^{app}$ (µM·s$^{-1}$) | $K_M^{app}$ (µM) | $k_{cat}^{app}$ (s$^{-1}$) | $k_{cat}^{app}/K_M^{app}$ (µM$^{-1}$·s$^{-1}$) |
| WT | 0.025±0.001 | 3.3±0.3 | 112±3 | 34±2 |
| AcK386 | 0.022±0.002 | 5±1 | 94±7 | 19±4 |
| HibK386 | 0.011±0.002 | 6±1 | 51±10 | 9±3 |
